# Experimental Evolution Reveals Limited Adaptation in Microbiota-less *Drosophila melanogaster*

**DOI:** 10.1101/2024.10.25.620247

**Authors:** Akshay Malwade, Anish Koner, B. Vibishan, Chinmay Gadkari, Sarthak Saini, Vinayak Khodake, Sutirth Dey

**Affiliations:** Indian Institute of Science Education and Research (IISER) Pune, Dr. Homi Bhabha Road, Pashan, Pune, Maharashtra, 411008, India

**Keywords:** experimental evolution, *Drosophila melanogaster*, gut microbiome, life history, evolutionary addiction, hygiene hypothesis

## Abstract

In single-generation experiments, microbiome removal often affects multiple host traits. However, relatively little is understood about the long-term implications of living in a microbe-free environment. To address this question, we evolved four replicate laboratory populations of *Drosophila melanogaster* without microbes and compared their traits with corresponding control populations that were reared with microbiota. This comparison was done under two assay environments: with-microbes and microbe-free. Prior single-generation experiments on our populations had revealed that microbiota removal significantly affects multiple traits, which suggested that the flies were under strong selection pressure to adapt to the absence of microbes. However, contrary to our expectations, the selected populations underwent very modest changes even after 54-57 generations of selection. Moreover, the magnitude of change in trait values across the with- and without- microbe environments was less for the selected populations than for the controls. RNA-Seq on one of the evolutionary replicates revealed that compared to the control, some anti-microbial peptides (AMPs) in the selected population were up-regulated, while several heat shock proteins (HSPs) were down-regulated. These results suggest that robust host-microbiome integrations on short timescales can nevertheless be labile on longer timescales. We situate these results in the context of the “evolutionary addiction hypothesis” and the “hygiene hypothesis”.

## Introduction

For millions of years, multicellular organisms have lived on this planet in conjunction with their associated microbial communities. Yet, the evolutionary dynamics of the symbiotic relationships between the hosts and their microbiota remain poorly understood. The hologenome theory of evolution, proposed nearly twenty years ago, suggests that the host and its associated microbiota—collectively termed the holobiont—can be considered a unified entity subject to natural selection (Rosenberg & Zilber-Rosenberg, 2018; Zilber-Rosenberg & Rosenberg, 2008). Since its inception, this theory has generated considerable interest and controversies within the scientific community (Bordenstein & Theis, 2015; Douglas & Werren, 2016; Madhusoodanan, 2019; Moran & Sloan, 2015; Morris, 2018). For example, critics have argued that the identity of the host-associated microbiota is often so dynamic (often changing within the host’s lifetime (Bana & Cabreiro, 2019)) that the holobiont cannot be considered a well-defined evolutionary unit. Others have pointed out that the interactions between a host and various components of its microbiota can range from beneficial to antagonistic (and everything in between), which leads to variation in selective pressures among constituents of host-microbiome assemblages (Douglas & Werren, 2016). This makes it difficult to consider the holobiont as a meaningful evolutionary unit. Despite the above-mentioned controversies about the status of the holobiont as a unit of selection/evolution, it is generally accepted that microbiota can play a significant role in a host’s evolution in multiple ways (Gilbert et al., 2015; Henry et al., 2021; Kolodny et al., 2020; Macke et al., 2017).

Theoretical studies have shown that under the conditions of strong vertical selection and short generation time, hosts can get a fitness advantage from symbiont microbes, even if the microbes pay costs for such an advantage (Vliet & Doebeli, 2019). It has been argued that the microbiome can alter the mean and the variance of the host’s phenotypic distribution, which in turn can enable the latter to explore a novel fitness landscape (Henry et al., 2021). These authors also suggest that the variance contributed by microbes can alter the heritability of the traits, thus affecting the host’s response to selection.

Empirical studies show that microbes can affect various facets of the biology of their hosts (reviewed in (Rosenberg, 2021)), including physiology (Grenier & Leulier, 2020), life history (Macke et al., 2017), behavior (Vuong et al., 2017), and immunity (Belkaid & Hand, 2014). Consequently, when the impact of the symbiont-mediated modulation is substantial, microbes are expected to alter the host’s fitness significantly (Ma et al., 2023; Rosenberg & Zilber-Rosenberg, 2016), thereby modulating their evolution.

Microbiota can confer rapid adaptation to the hosts by expanding the host’s niche (Borges, 2017; Hoang et al., 2021; Koide, 2023; Kopac & Klassen, 2016). For example, a study showed that stinkbugs can gain insecticide resistance within a single generation by acquiring the bacterial strain of *Burkholderia* that can degrade the insecticide fenitrothion (Kikuchi et al., 2012). This study also showed that along with insecticidal resistance, stinkbugs gained fitness advantages such as higher survival and larger body size that could potentially help this symbiosis spread in the population. Furthermore, it is known that symbiotic microbes can be an important source of phenotypic plasticity for their host (Alberdi et al., 2016), which can serve as an initial step in the host’s adaptation to novel environments.

Given the diverse roles that microbiota can play in the evolution of their hosts and the fact that almost all organisms have been co-habiting with microbes for millions of years, it is natural to ask what will happen to organisms if they are forced to live without microbes for long periods of time. We already know that in some species, including caterpillars of tobacco hornworms (Hammer et al., 2017) and butterflies (Phalnikar et al., 2019), removing the microbiota has little or no effects. However, in the vast majority of species, removing the microbiota leads to prominent physiological and behavioral effects (Rosenberg, 2021). If subjected to prolonged microbiota absence, will such species fail to grow beyond a few generations, or will they adapt? If the latter, will the adaptation be fast or slow? Will their ability to adapt to other stresses be affected by the prolonged absence of the microbes?

While the above questions are of interest to evolutionary biologists, there are more pragmatic anthropocentric reasons for understanding the long-term impact of microbiota on host evolution. In modern human societies, the overuse of antibiotics and a hygienic lifestyle, along with other factors, could lead to the disappearance of beneficial microbes (Blaser & Falkow, 2009; Noverr & Huffnagle, 2004). It has been hypothesized that such loss of ancestral microbes (e.g., *H. pylori*) can increase the risk factors for conditions such as gastroesophageal reflux disease (GERD) and childhood asthma (Blaser, 2008; Blaser & Falkow, 2009). Even the microbiota of many naturally occurring species might have been altered due to increased antibiotic load in the environment (Baquero et al., 2019; Larsson & Flach, 2022) and micro-environmental alterations brought by climate change (Williams et al., 2023). Therefore, understanding the evolutionary implications of altering or removing the microbiota on the hosts has emerged as a key question in biology (Macke et al. 2017; Henry et al. 2021).

Here, we used experimental evolution on *D. melanogaster* populations to understand how a host evolves in the absence of its microbiota. In this species, removing microbiota for a single generation can lead to developmental delay, lower body size (Shin et al., 2011; Storelli et al., 2011), reduced female fecundity (Elgart et al., 2016; Gnainsky et al., 2021; Suyama et al., 2023), and changes in behavioral traits such as diet-induced mating preference (Sharon et al., 2010), foraging (Wong et al. 2017), aggression (Jia et al., 2021), and locomotor activity (Schretter et al., 2018). Thus, one expects that in *Drosophila*, the removal of the microbiota would affect the hosts’ fitness in multiple ways, potentially leading to a strong, multi-directional selection pressure. This makes *Drosophila* a well-suited system for studying this question.

We kept four large (∼2400 individuals) outbred laboratory populations of *D. melanogaster* microbe-free using bleach (labeled ‘MBL’ populations, for ‘<u>M</u>icro<u>b</u>iota-<u>L</u>ess’) for more than 50 generations. This involved bleaching *D. melanogaster* eggs every generation and rearing them in sterile food in a sterile environment. Each of these four selected populations had an ancestry-matched control (labeled ‘MB’ populations) where the microbiome was reconstituted from a common source pool just after the bleach treatment. For this reconstitution, microbes were sampled from the environment of the flies via fecal microbiome transfer (Supplementary Figure S9 shows the composition of the ancestral microbiome that is dominated by *Acetobacter* and *Lactobacillus* spp.). We compared the selected MBL and control MB populations in two assay environments: (a) in the presence of microbes (labeled “with-microbes” environment) and (b) in the absence of microbes (labeled “microbe-free” environment). We hypothesized that the host would face substantial selection pressure in the absence of microbes. Therefore, we expected a rapid evolution of the host to maintain the homeostasis seen in the presence of microbes in our ancestral baseline populations. We found that even after surviving for 54 generations without microbes, contrary to expectations, there were only modest adaptations in the selected populations. Compared to the controls, the trait values of the selected populations were found to be less affected by the absence of microbes in the assay environment.

RNA-Seq on one of the evolutionary replicates suggested that a cluster of anti-microbial peptides (AMPs) was up-regulated, and another cluster of heat shock proteins (HSPs) was down-regulated in the MBLs. Finally, we discuss the potential implications of these results in terms of existing theories about host-microbiota evolutionary relationships.

## Methods

### Details of the experimental evolution lines

The experimental evolution lines were derived from the four ancestral baseline populations named NDB_1-4_ (at their generation 21 in the lab). The details about the NDB_1-4_ are provided in Supplementary Text S1.1. Two populations were derived from each ancestral NDB line: MBL (“<u>M</u>icro<u>b</u>iota-<u>L</u>ess,” selected population that is without the microbiome) and MB (corresponding ancestry-matched control population with microbiome) (Supplementary Figure S1). For example, NDB_1_ gave rise to MBL_1_ and MB_1_, NDB_2_ to MBL_2_ and MB_2_, and so on. Thus, populations with the same subscript (e.g., MBL_2_ and MB_2_) share a common ancestry and were always assayed together. Each population in this setup is maintained at a size of ∼2400 adult flies in plexiglass cages with an additional microbiome removal protocol in each generation, as described in the next section. NDB_1_ served as the common source of the microbiome for all the MBs (for the rationale of this step, see microbiome reconstitution section). All populations were maintained at 25°C and constant light conditions.

### Experimental evolution protocol employed each generation

MBLs were handled aseptically inside a Level II biosafety cabinet (Microfilt, India). Any material that came in contact with MBLs for their maintenance was either autoclaved, filtered through a 0.2-micron filter, or surface-sterilized with 70% ethanol.

Every generation, on day 12 after egg collection, each population of MBLs and MBs (a total of eight populations) was provided with a moist paste of autoclaved yeast to boost their egg output. This paste also contained a few drops of acetic acid so that the flies were attracted to yeast. A fresh food plate with an exposed vertical food surface was provided for 12-16 hours to these populations at the end of day 13 to collect eggs for the next generation. On day 14, these eggs were collected using a moist paintbrush and transferred to cell strainers (TCP026, HiMedia) for egg dechorionation (Supplementary Figure S2A). The cell strainer was then placed into one of the wells of a 6-well plate. Based on known protocols to rear microbe-free *Drosophila* eggs (Kietz et al., 2018; Koyle et al., 2016; Ridley et al., 2013), these eggs were washed with 2% bleach for 2 min to remove the chorion layer and any associated microbes (Supplementary Figure S2B). The cell strainer with dechorionated eggs was transferred to another well of the 6-well plate for another quick wash with 2% bleach to kill any residual microbes. Next, the cell strainer with eggs was transferred back-to-back to two fresh wells of the 6-well plate for two successive washes with autoclaved dH_2_O to remove bleach. For the MBLs, ∼400 eggs were transferred to translucent plastic milk bottles (Laxbro Inc., Catalog No. FLBT-20) with ∼50-60 ml autoclaved solid banana-jaggery media (Supplementary Figure S2C). To make MBs, dechorionated eggs were reconstituted with the native microbiome of NDB_1_ (see next section) before being transferred to the food bottles (Supplementary Figure S2C).

The presence of microbes in MBs and their absence in MBLs is shown by PCR using 16S rRNA universal bacterial primers in Supplementary Figure S5 as described in Supplementary Text S3.1 and S3.2.

### Microbiome reconstitution

For long-term experimental evolution, we chose microbiome reconstitution to create flies with microbes. Supplementary Figure S3 illustrates the reconstitution protocol in detail. Microbiome from the ancestral NDB_1_ was used in reconstitution for all the MBs. This ensured that there were no differences across the four MB populations in terms of the microbiota that they harbored. Several distinct sources of the microbiome (say, every MB_i_ getting their microbiome from their ancestral NDB_i_) would have meant a greater chance of differential microbial association, which we sought to avoid.

Eight fly vials with non-autoclaved standard banana-jaggery food were set to house the flies for 24 hrs. These flies were collected on day 13 after egg collection. 50 flies with an equal male-to-female ratio were transferred into each vial using mild CO_2_ anesthesia. After 24 hours, the flies were discarded, and 1 ml of autoclaved water was added to each vial to extract the microbiome. Vials were swirled with water till no visible fecal spots were seen on the walls. This method of sampling of microbes from the frass of flies housed in vials has been used earlier in the literature (Chandler et al., 2022). The microbe-free eggs, after the bleach treatment, were washed with autoclaved water twice and then dipped twice into this microbiome slurry to seed the source microbiome on the eggs. These eggs were then gently transferred to *Drosophila* vials or bottles with autoclaved food to generate flies with reconstituted microbiomes derived from NDB_1_.

### Common-garden assay environments for the phenotypic assessment

We explain the assay environments using the MB_1_-MBL_1_ (Figure 1) as an example. However, the same is true for the rest of the three groups of MB-MBL as well. MB_1_ received microbiome from NDB_1_ every generation.

**Figure 1.**
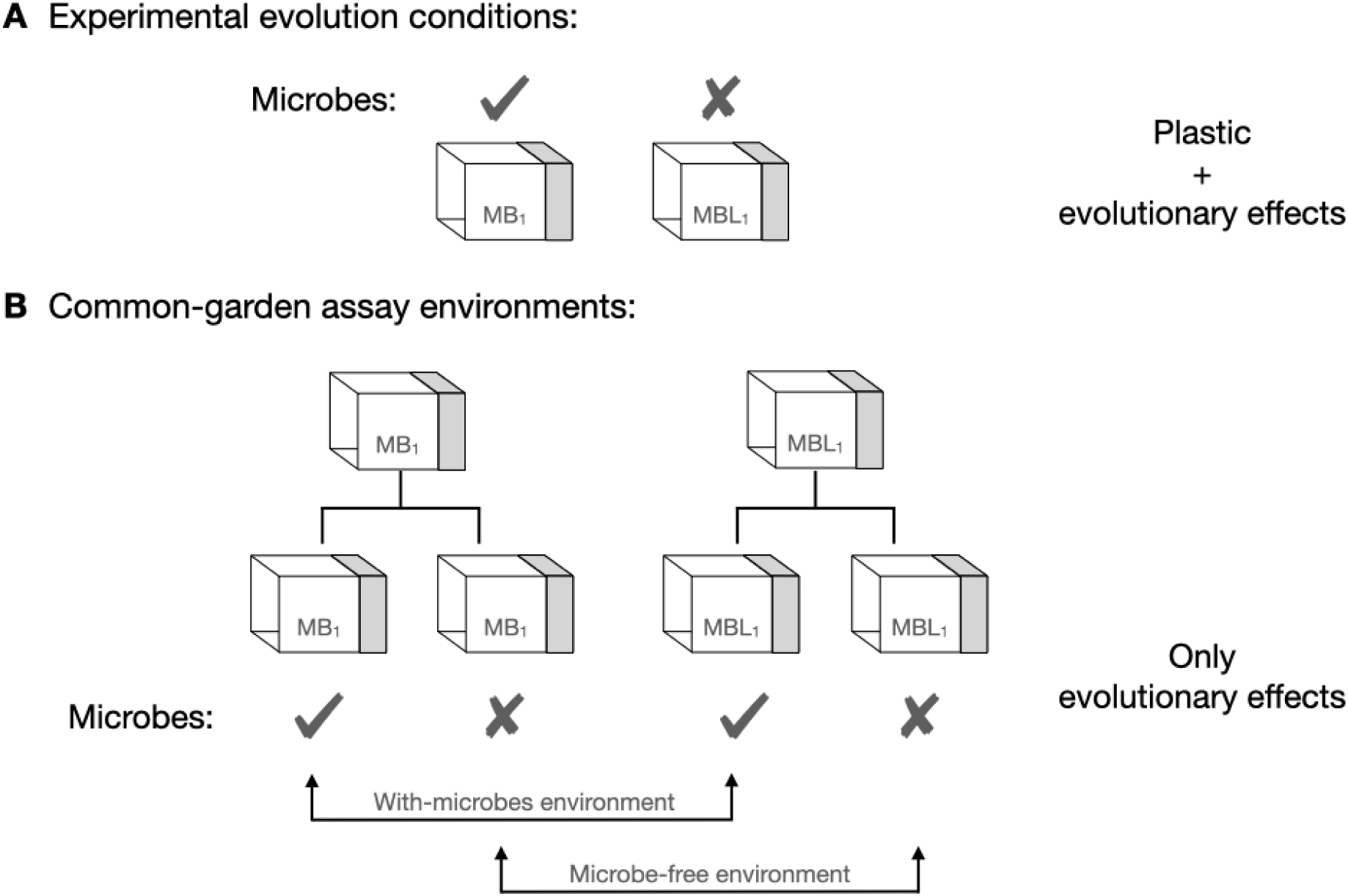
Illustration of environments in which experimental evolution was carried out and the environments in which phenotypic assays are carried out.

MBL_1_ is maintained without any microbes. If we directly compare the phenotypes of MB_1_-MBL_1_ in their experimental evolution conditions (i.e., MB with microbes and MBL without microbes) (Figure 1A), any observed differences between MB/MBL lines can be due to their evolutionary history (with/without microbes) or the assay environment (with/without microbes) or both. Therefore, we needed to provide a common assay condition for the phenotypic comparison of MB_1_-MBL_1_. We performed a phenotypic comparison between MB_1_-MBL_1_ both in the native environment of MB_1_, which is with-microbes environment, as well as in the native environment of MBL_1_, which is microbe-free (Figure 1B). For the with-microbe environment, MB_1_ as well as MBL_1_ were maintained in this common-garden environment in the presence of microbes for a generation. Assays were then performed on F1 flies (F1 flies generated from the selected and control flies that underwent 17-20 or 54-57 generations of selection) from the with-microbe environment that were raised as described in Supplementary Text S2 and Supplementary Figure S4. For a microbe-free environment, MB_1_ and MBL_1_ were maintained in this common-garden environment in the absence of microbes for a generation. Microbe-free F1 flies on which we performed the assays were raised as described in Supplementary text S2 and Supplementary Figure S4. Matching of environments via common-garden rearing for a generation minimizes non-genetic effects and allows us to attribute any observed differences in phenotype to the evolutionary effects (instead of effects due to short-term plastic changes) in which we are primarily interested. The assay design aims to ensure that only evolutionary effects contribute to the selected vs. control comparison.

The detailed validation of these assay conditions is reported in Supplementary Text S5.1-S5.2.

### Assays

We measured various traits in the selected and control populations twice, first in generations 17-20 and then in generations 54-57 (the experimental design details of these two assessments are given in Supplementary Text S4.1 and S4.2, respectively).

For all the following assays, eggs were dechorionated as explained in a previous section (Supplementary Figure S2) and transferred into plastic vials with ∼ 6 ml of food at a pre-determined density (∼30 eggs per vial unless stated otherwise). All the assays were carried out in autoclaved banana-jaggery food.

#### Egg-to-adult developmental time and Percentage survival to adulthood

A total of ten replicate vials were used per treatment. For each vial, we counted precisely 30 eggs under a microscope using a moist paintbrush and transferred them to a thin strip of solidified 1.3% non-nutritive agar. This piece of agar with exactly 30 eggs was then transferred to the food vial with autoclaved banana-jaggery food. The vials were kept in an incubator at 25°C under constant light conditions. When the flies were about to eclose, the vials were monitored at regular intervals of 2 hours (generations 17-20) or 6 hours (generations 54-57). Every two hours, the males and females were sex separated under mild CO_2_ anesthesia, and their numbers were noted. When there were no further eclosions in a vial, we counted the total number of eclosed flies and divided that by 30 (i.e., the number of eggs put in the vial) to compute the egg-to-adult survivorship.

#### Dry body weight

On day 12 after egg collection, the adult flies were anesthetized (using CO_2_) and sorted by sex into batches of 20 individuals per sex per tube into 1.5 ml micro-centrifuge tubes. These flies were then flash-frozen using liquid nitrogen and, if required, stored at -80^0^C till weighing. The flies were then air dried at 60^0^C for 72 hrs and weighed on an analytical weighing balance (ME104, Mettler Toledo, least count = 0.1 mg). First, the total weight of (micro-centrifuge tube + flies) was recorded. Next, flies were removed from the tubes with a dry paintbrush, and the weight of the empty micro-centrifuge tube was recorded. The difference between these two gave us the body weight of 20 flies. This number was divided by 20 to get the average dry body weight per fly. When flies were removed, we counted these flies to double-check if every micro-centrifuge tube had 20 flies. If not, then the total weight was divided by that count. A total of ten micro-centrifuge tubes were weighed per treatment per sex. Thus, 200 flies per treatment per sex were sampled.

#### Female fecundity

On day 12 after egg collection, flies were transferred to fresh food vials. On day 13 (which is the day on which eggs are collected during the selection process), a male-female pair was aspirated (to avoid CO_2_ anesthesia) into a 50 ml centrifuge tube with several holes in the wall for ventilation. The falcon tube had a food cup stuck to the inner surface of its cap to provide a surface for egg-laying. The centrifuge tubes were kept for 12 hours in an incubator maintained at 25^0^C and constant light, after which the total number of eggs laid on the food cup was counted under a microscope. A total of 50 replicates per treatment were set for this assay.

#### Desiccation resistance

Ten 12-day old adult flies of a given sex were introduced into an empty plastic vial (Height: 3.5 inches x Diameter: 1 inch) using aspiration to avoid CO_2_ anesthesia. The flies had no access to food or water. The vials were plugged with fresh cotton plugs and checked manually every two hours for any deaths till all the flies were dead. At each time point, the number of live flies was recorded (total flies to start with - count of total dead flies). The average time to death due to desiccation was calculated for each vial. Ten such replicate vials were set per treatment per sex.

#### Locomotor activity

The flies’ locomotor activity was measured using the *Drosophila* Activity Monitoring (DAM) system (Trikinetics Inc). Using an aspirator, a single adult fly (day 12 after egg collection) was introduced into a glass tube (Length: 6.5 cm, Diameter: 5 mm). The locomotor activity for a given fly is the average number of times the fly crosses the infrared beam running through the middle of the glass tube per hour, recorded over six hours. Approximately 32 flies were used per treatment per sex. Data from flies that were seen to be dead at the end of the six-hour recording period were excluded from the analysis. The first 15 minutes of activity were excluded from the analysis as the time taken for the flies to get acclimatized to the glass tubes.

#### Mating latency and mating time

The mating assay was performed on virgin flies. After eclosion (typically around day eight after egg collection), male and female flies were separated under mild CO_2_ anesthesia into vials with fly food. To ensure that the eclosed flies were virgins, we collected the flies every six hours after the first eclosion (Demerec, 1967). Males and females were held separately at the density of ten flies per vial till the setting up of the mating experiment on day 12 after egg collection.

On the day of the experiment, a female was introduced into a fresh plastic vial using an aspirator, followed by the introduction of a male. The floor of the vials was already covered with solid 1.3% non-nutritive agar to avoid desiccation of the flies. The time at which the male fly was introduced (T1), the time at which mating started (T2), and the time at which mating ended (T3) were noted through live manual observation. The mating latency was calculated as (T2-T1), and the mating time was calculated as (T3-T2). A total of 40 replicates per treatment were used for this assay.

#### Robustness to the presence/absence of microbes

In this study, we looked at the effects of the presence/absence of microbiota across multiple traits. To get an overall picture from this data, we needed to look at a combined measure of the magnitude of the effect of microbiota removal on the trait values. We call this measure the robustness to microbiota removal (or simply, robustness) and conceptualize it as the difference in the average trait value between the with-microbe and the microbe-free conditions. Higher robustness refers to a lower average change in the trait value due to microbiota removal and vice versa. For this, one needs a dimensionless (or a normalized) measure of the trait value, as the various traits measured here (e.g., mg of bodyweight, no. eggs laid, percent survival) have different units. We used two normalized/dimensionless measures that fit the above criteria and then compared the mean robustness for the MB-MBL populations.

The first measure was the effect size (Cohen’s d) for the MB vs. MBL populations across the two environments: with- and without microbes (Cohen, 2013). In other words, for every trait in each population, we computed the effect sizes of the difference between the trait values in the with-microbe and the microbe-free environments. Thus, for example, for the development time of MBs, we computed the effect size of the difference between the mean development time over the four populations in the with-microbe environment and the microbe-free environment. We reasoned that whichever populations (MBs or MBLs) showed a greater effect size due to microbiome removal for a particular trait exhibited lower robustness for that trait across the with-microbe and microbe-free environments. It should be noted here that these across-environment effect sizes are different from those provided in Tables 1-2 and Supplementary Tables S5-S6. Supplementary Texts S4.1 and S4.2 discuss the experimental design used to calculate these plastic effect changes and why such analysis could not be done in Gen 17-20.

**Table 1.** Summary of all assays in the <u>with-microbes environment (Gen 54-57).</u> T refers to the grand mean of the MBLs, while C refers to the grand mean of the MBs.

|  | Assay | Test statistic | P-value | % change (T - C)/C | Cohen's d | Effect size* | Sample size<sup>§</sup> |
| --- | --- | --- | --- | --- | --- | --- | --- |
| 1 | Dry body weight (Male) | $t_3 = 0.32$ | 0.7723 | -0.6 | 0.07 | Small | 10 |
| 2 | Dry body weight (Female) | $t_3 = 0.82$ | 0.4726 | -1.8 | 0.17 | Small | 10 |
| 3 | Development time (Males) | $t_3 = 2.62$ | 0.0788 | 0.4 | 0.17 | Small | 9 to 10 |
| 4 | Development time (Females) | $t_3 = 0.91$ | 0.4299 | 0.2 | 0.06 | Small | 10 |
| 5 | Survival to adulthood | $t_3 = 2.31$ | 0.1044 | -2.7 | 1.81 | Large | 9 to 10 |
| 6 | Fecundity (Females) | $t_3 = 1.75$ | 0.1791 | 26.0 | 1.67 | Large | 50 |
| 7 | Desiccation resistance (Males) | $t_3 = 3.16$ | 0.0510 | -18.2 | 1.62 | Large | 10 |
| 8 | Desiccation resistance (Females) | $t_3 = 2.45$ | 0.0916 | -6.1 | 0.89 | Large | 10 |
| 9 | Locomotor activity (Males) | $t_3 = 0.50$ | 0.6498 | 6.4 | 0.37 | Small | 30 to 32 |
| 10 | Locomotor activity (Females) | $t_3 = 0.10$ | 0.9260 | 1.1 | 0.07 | Small | 31 to 32 |
| 11 | Mating Latency | $t_3 = 0.72$ | 0.5211 | 11.1 | 0.44 | Small | 34 to 40 |
| 12 | Mating time | $t_3 = 0.15$ | 0.8928 | -0.4 | 0.04 | Small | 34 to 40 |

**Table 2.**
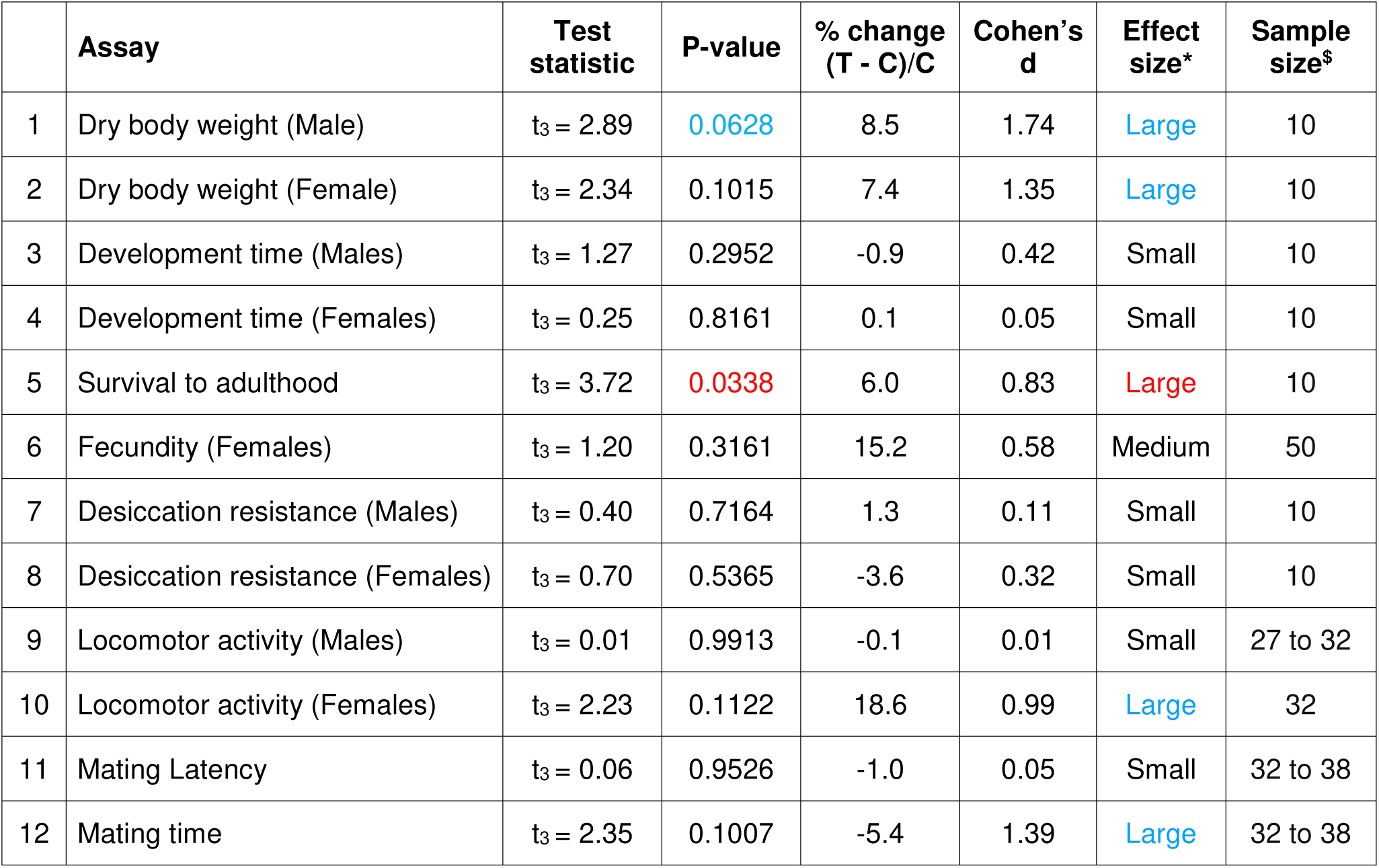
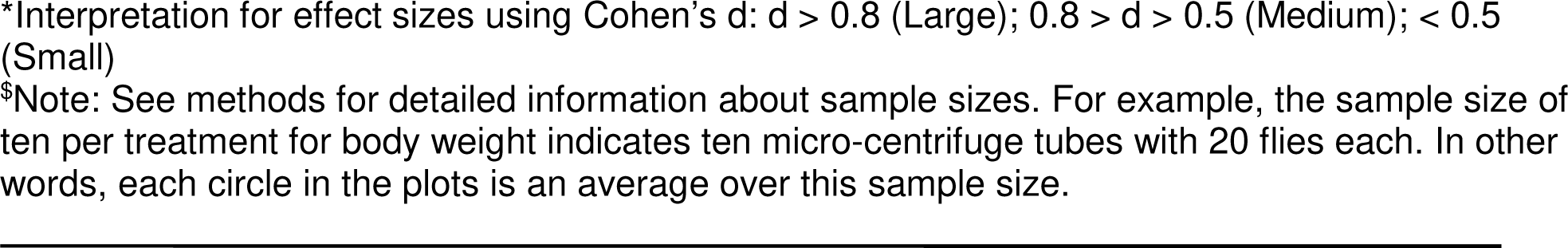
Summary of all assays in the <u>microbe-free environment (Gen 54-57)</u>. T refers to the grand mean of the MBLs, while C refers to the grand mean of the MBs.

|  | Assay | Test statistic | P-value | % change (T - C)/C | Cohen's d | Effect size* | Sample size<sup>§</sup> |
| --- | --- | --- | --- | --- | --- | --- | --- |
| 1 | Dry body weight (Male) | $t_3 = 2.89$ | 0.0628 | 8.5 | 1.74 | Large | 10 |
| 2 | Dry body weight (Female) | $t_3 = 2.34$ | 0.1015 | 7.4 | 1.35 | Large | 10 |
| 3 | Development time (Males) | $t_3 = 1.27$ | 0.2952 | -0.9 | 0.42 | Small | 10 |
| 4 | Development time (Females) | $t_3 = 0.25$ | 0.8161 | 0.1 | 0.05 | Small | 10 |
| 5 | Survival to adulthood | $t_3 = 3.72$ | 0.0338 | 6.0 | 0.83 | Large | 10 |
| 6 | Fecundity (Females) | $t_3 = 1.20$ | 0.3161 | 15.2 | 0.58 | Medium | 50 |
| 7 | Desiccation resistance (Males) | $t_3 = 0.40$ | 0.7164 | 1.3 | 0.11 | Small | 10 |
| 8 | Desiccation resistance (Females) | $t_3 = 0.70$ | 0.5365 | -3.6 | 0.32 | Small | 10 |
| 9 | Locomotor activity (Males) | $t_3 = 0.01$ | 0.9913 | -0.1 | 0.01 | Small | 27 to 32 |
| 10 | Locomotor activity (Females) | $t_3 = 2.23$ | 0.1122 | 18.6 | 0.99 | Large | 32 |
| 11 | Mating Latency | $t_3 = 0.06$ | 0.9526 | -1.0 | 0.05 | Small | 32 to 38 |
| 12 | Mating time | $t_3 = 2.35$ | 0.1007 | -5.4 | 1.39 | Large | 32 to 38 |

\*Interpretation for effect sizes using Cohen's d: $d > 0.8$ (Large); $0.8 > d > 0.5$ (Medium); $< 0.5$ (Small)
§Note: See methods for detailed information about sample sizes. For example, the sample size of ten per treatment for body weight indicates ten micro-centrifuge tubes with 20 flies each. In other words, each circle in the plots is an average over this sample size.

The second measure that we used for robustness was the percentage change in the magnitude of the traits across environments.

### RNA isolation, RNA-Seq, and analysis

A subset of flies from the phenotypic assays (at generation 54-57) were flash-frozen in liquid nitrogen and stored at -80^0^C. The frozen flies from the MB_1_ and MBL_1_ populations were sent to a commercial vendor (miBiome Therapeutics LLP, Mumbai, India) for RNA-Seq, starting with total RNA isolation. Due to logistical constraints, RNA-Seq was done only on female flies. The details of RNA-Seq are provided in Supplementary Text S1.6.

### Statistical analysis

The selected populations (MBL_1-4_) were compared with their corresponding controls (MB_1-4_) in two types of environments (with and without microbiota). The analysis for these two environments was done separately. In each environment, means of MB_1-4_ were compared against means of MBL_1-4_ using a paired t-test performed in GraphPad Prism (v10.2.1). We compared the robustness of MBs and MBLs to microbiome removal (i.e., Cohen’s *d* and percentage change across environments) using separate paired t-tests.

For interpreting effect sizes using Cohen’s d, the following criteria were followed: d > 0.8 (high); 0.8 > d > 0.5 (medium); d < 0.5 (low) (Cohen, 2013). The t-tests were performed, and the graphs were plotted in GraphPad Prism (v10.2.1). The figures drawn using BioRender are labeled as such in their description.

In the Results section, along with the trends that are statistically significant (i.e., P < 0.05), we have labeled results where 0.05 < P < 0.1 and Cohen’s d ≥ 0.8 (i.e., large effect size) as trends. This is highlighted in blue in Tables 1-2 and Supplementary Tables S5-S6, which provide detailed information about the statistical comparisons for all the assays.

## Results

### First phenotypic assessment of experimental evolution lines at host generations 17-20

The results for generations 17-20 are provided in Supplementary Text S6. At this point, we observed that barring mating time (which was significantly shorter in MBLs), none of the other traits differed between the MBs and the MBLs (Supplementary Tables S5 and S6).

### Second phenotypic assessment of experimental evolution lines at host generations 54-57

Tables 1 and 2 provide detailed information on the statistical comparison between MB_1-4_ and MBL_1-4_ for the with-microbe and microbe-free assay environments, respectively.

Comparing these tables with the corresponding ones for generation 17 (Supplementary Tables S5 and S6) shows that there were a larger number of cases in the later generation where the effect sizes were medium or large. This could be a signature of evolutionary divergence between the MB and MBL populations.

#### Male traits in host generations 54-57

There was no difference between the body weight of the male MBL_1-4_ and MB_1-4_ flies in the with-microbe assay environment (Table 1, Figure 2A-left). In the microbe-free environment, the body weight of the male MBL_1-4_ was more than their corresponding controls MB_1-4_ with a large effect size even though the P-value was marginally greater than 0.05 (Table 2, Figure 2A-right).

**Figure 2.**
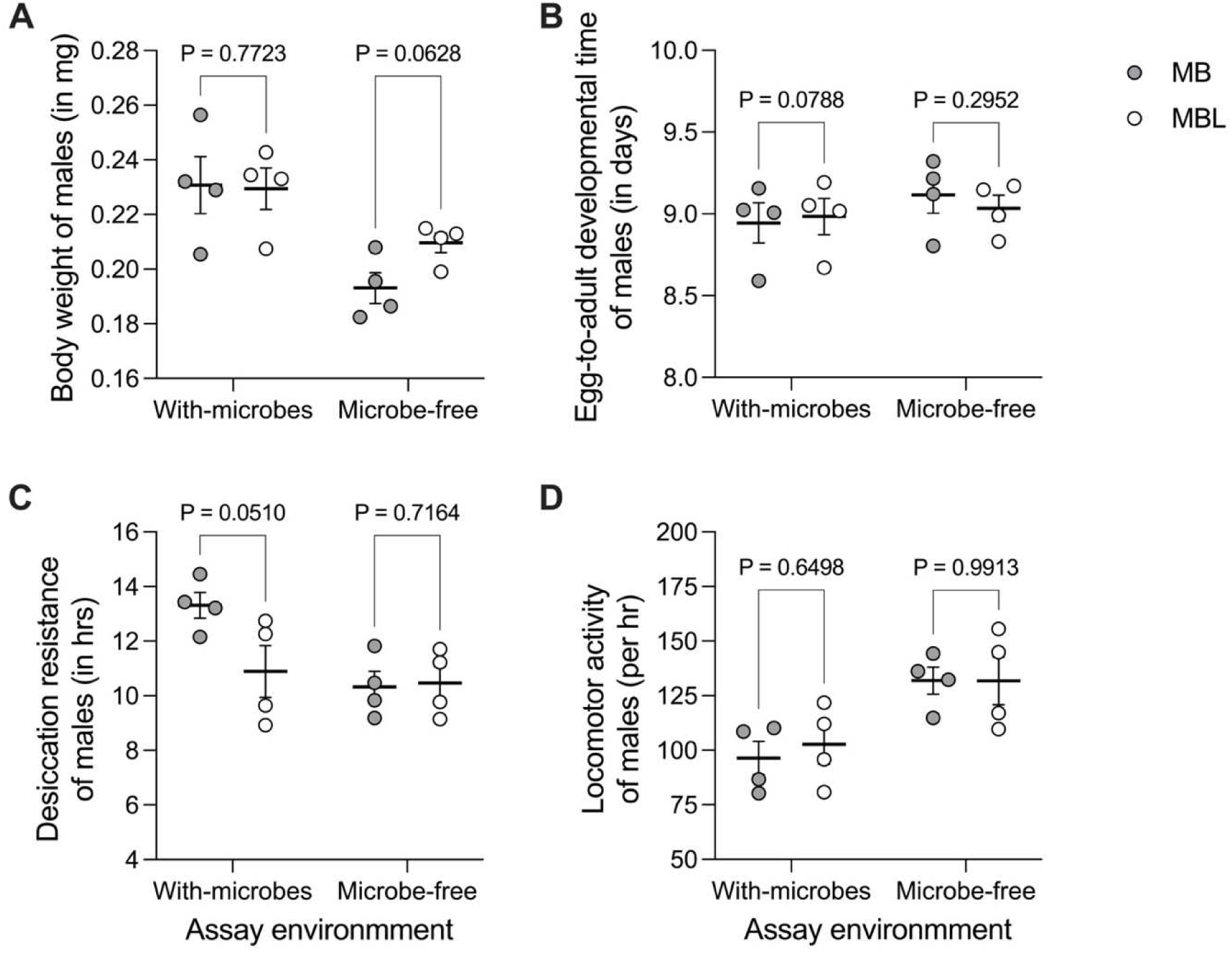
Male traits assessed in Gen 54-57 in with-microbe and microbe-free environments. (A) Body weight. (B) Egg-to-adult development time. (C) Desiccation resistance. (D) Locomotor activity using DAM system. In each assay environment, four filled circles are means for each of the four control populations MB_1-4_ and, similarly, four empty circles are means for each of the four selected populations MBL_1-4_. The black horizontal lines represent the grand mean over four population means and the error bars represent the SEM. For each host phenotype, the grand mean over MB_1-4_ was compared against the grand mean over MBL_1-4_ using a paired t-test.

The egg-to-adult development time was largely unaffected by the multi-generation deprivation of microbes, as seen in both assay environments (Tables 1 and 2, Figure 2B).

MBL_1-4_ males had lower desiccation resistance than MB_1-4_ in the with-microbe environment with a large effect size, although the P-value was marginally greater than 0.05 (Table 1, Figure 2C-left). However, no difference was observed in the microbe-free environment (Table 2, Figure 2C-right).

Likewise, MB_1-4_ and MBL_1-4_ showed similar levels of locomotor activity in both environments (Tables 1 and 2, Figure 2D).

#### Female traits in host generations 54-57

The body weights of the MB_1-4_ and MBL_1-4_ females were similar in the with-microbes regime (Table 1, Figure 3A-left) but showed a marginal increase for MBLs in the no-microbe regime, which was not statistically significant (Table 2), despite a large effect size (d = 1.35, Figure 3A-right).

**Figure 3.**
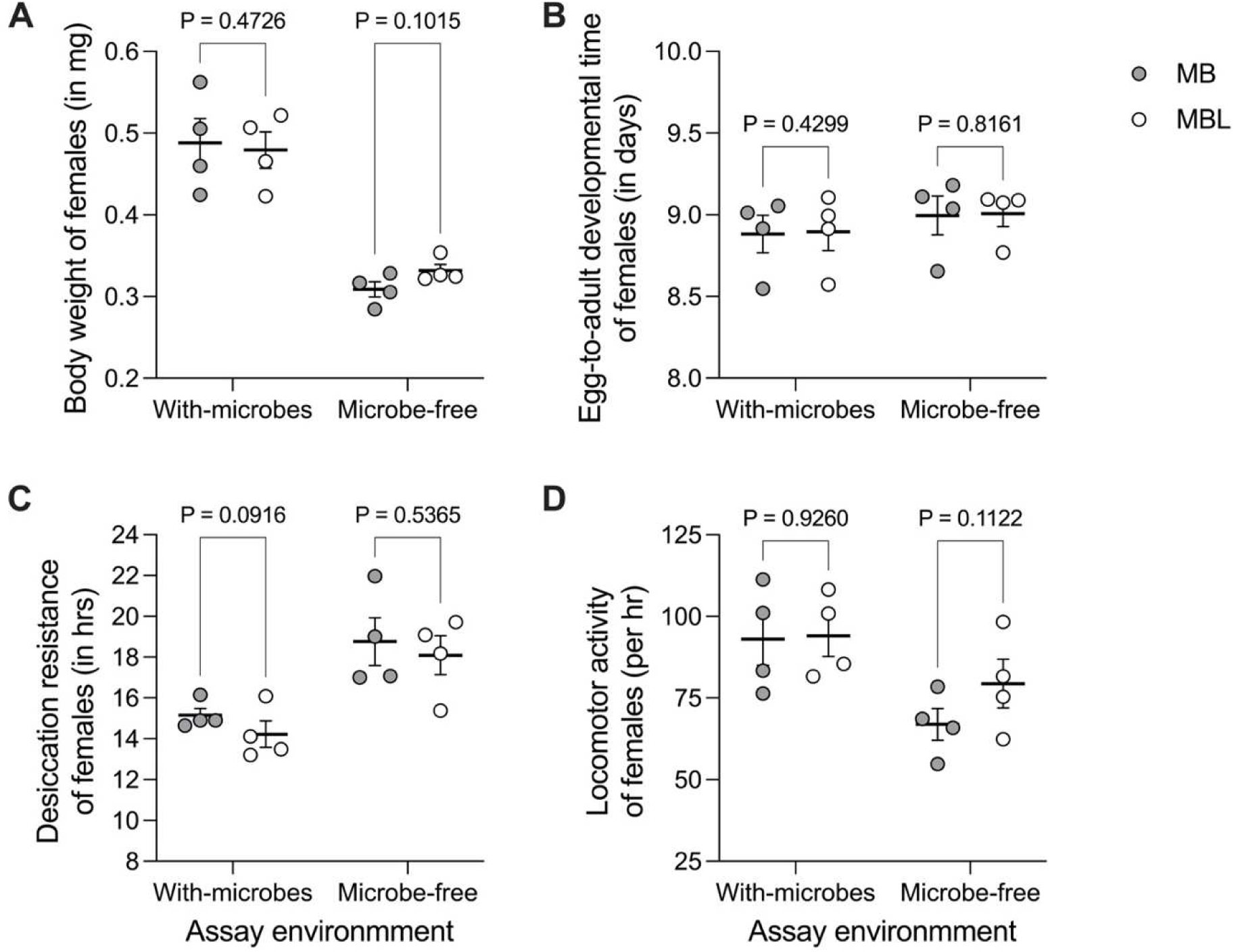
Female traits assessed in Gen 54-57 in with-microbe and microbe-free environments. (A) Body weight. (B) Egg-to-adult development time. (C) Desiccation resistance. (D) Locomotor activity using DAM system. In each assay environment, four filled circles are means for each of the four control populations MB_1-4_ and, similarly, four empty circles are means for each of the four selected populations MBL_1-4_. The black horizontal lines represent the grand mean over four population means and the error bars represent the SEM. For each host phenotype, the grand mean over MB_1-4_ was compared against the grand mean over MBL_1-4_ using a paired t-test.

Female egg-to-adult development time was similar for MB-MBLs after multiple generations without the microbes when assayed in both assay environments (Tables 1 and 2, Figure 3B).

MBL_1-4_ females had lower desiccation resistance in the with-microbe regime, which was not statistically significant (Table 1, Figure 3C-left). MBs and MBLs had similar desiccation resistance in the no-microbe regime (Table 2, Figure 3C-right).

MB_1-4_ and MBL_1-4_ females had similar locomotor activity levels in the with-microbes environment (Table 1, Figure 3D-left). In the microbe-free environment, there was a trend (i.e., large effect size, d = 0.89) towards MBL_1-4_ females being more active than the MB_1-4_ that was not statistically significant (Table 2, Figure 3D-right).

#### Pair traits in host generations 54-57

There was a trend (d = 1.81) of lower survival for MBL_1-4_ than MB_1-4_ in the with-microbe regime, but it was not statistically significant (Table 1, Figure 4A-left). In the microbe-free environment, MBL_1-4_ had better survival than MB_1-4_ (Table 2, Figure 4A-right).

**Figure 4.**
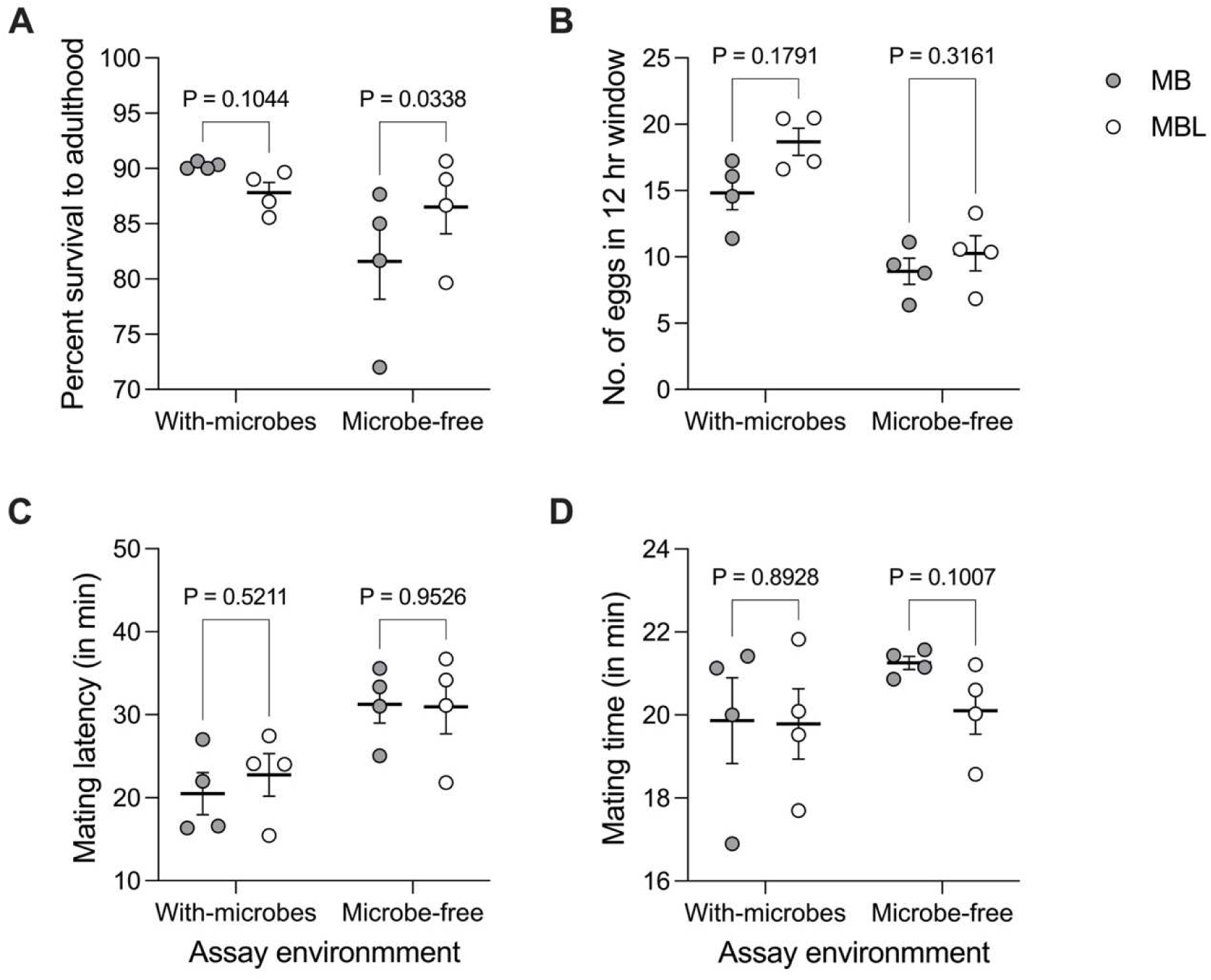
Pair traits assessed in Gen 54-57 in with-microbe and microbe-free environments. (A) Survival to adulthood. (B) Female fecundity. (C) Mating latency. (D) Mating time. In each assay environment, four filled circles are means for each of the four control populations MB_1-4_ and similarly, four empty circles are means for each of the four selected populations MBL_1-4_. The black horizontal lines represent the grand means over four population means and the error bars represent the SEM. For each host phenotype, the grand mean over MB_1-4_ was compared against the grand mean over MBL_1-4_ using a paired t-test.

There was a trend (d = 1.67) of MBL_1-4_ laying more eggs than the MB_1-4_ in the with-microbe regime, but this difference was not statistically significant (Table 1, Figure 4B-left). In the microbe-free environment, MBL_1-4_ had a trend of slightly higher fecundity, but it was not statistically significant either (Table 2, Figure 4B-right).

MBL_1-4_ did not differ from MB_1-4_ in the latency to start mating, both in the with-microbe and microbe-free regimes (Table 1 and 2, Figure 4C).

There was no difference in MB_1-4_ and MBL_1-4_ mating time in the with-microbe regime (Table 1, Figure 4D-left). In the microbe-free environment, there is a trend (d = 1.39) of shorter mating times for MBL_1-4_ than MB_1-4_, but this is not statistically significant (Table 2, Figure 4D-right).

### MBLs have greater robustness than the MBs across with-microbe and microbe-free environments (Gen 54-57 assays)

Figures 2 to 4 suggested that, compared to the MBs, the trait values of the MBLs changed relatively less across the two assay environments (i.e., with-microbes and microbe-free). In other words, the reaction norms for the various traits across these two environments are expected to be flatter for the MBLs than for the MBs. True to this observation, we found that the MBLs, on average, had smaller effect sizes than MBs (Paired t-test, t_11_ = 2.14, P = 0.0554, Cohen’s d = 0.49 (small), Figure 5A), hinting at lower magnitude of change (and hence greater robustness to microbiome removal) than the MBs across two assay environments. Moreover, MBLs had a lower percentage change in their trait values across two environments than MBs (Paired t-test, t_11_ = 2.99, P = 0.0123, Cohen’s d = 0.39 (small), Figure 5B).

**Figure 5.**
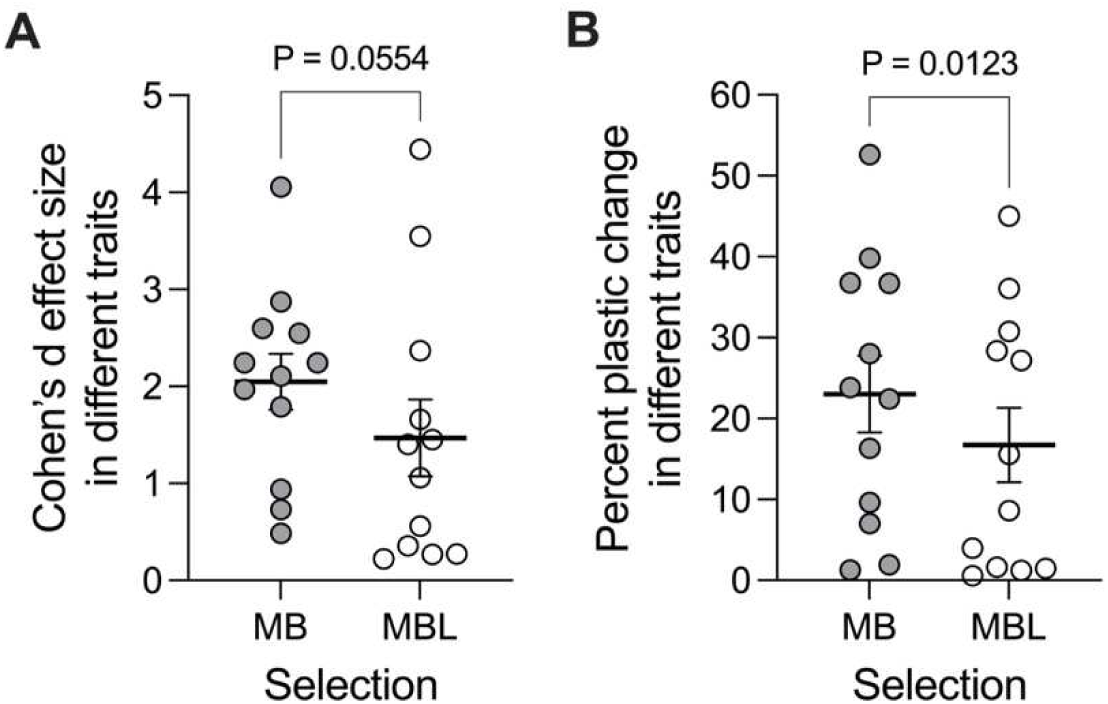
Robustness to microbiota removal. (A) Effect sizes and (B) Percentage changes. Means are indicated by black horizontal lines. The error bars represent SEM. The data is analyzed using a paired t-test. For percent change, Wilcoxon matched-pairs signed rank test (a non-parametric test) is also significant (W = -64.0, P = 0.0093).

Taken together, the MBLs had greater robustness (i.e., lesser effect of microbiota removal) than the MBs, even though the effect size was small for both the measures we employed.

### The big picture across both assessments: divergence in MB-MBLs with time?

In the previous section, we used effect sizes on phenotype differences across assay environments for MBs as well as MBLs to measure the robustness exhibited by these two populations. In this section, we look at how effect sizes have changed *for the evolutionary difference between MB-MBLs over time* from the first phenotypic assessment in generations 17-20 to the second assessment in generations 54-57 (Figure 6). These effect sizes are provided in Supplementary Tables S5-S6 for generations 17-20 and Tables 1-2 for generations 54-57.

**Figure 6.**
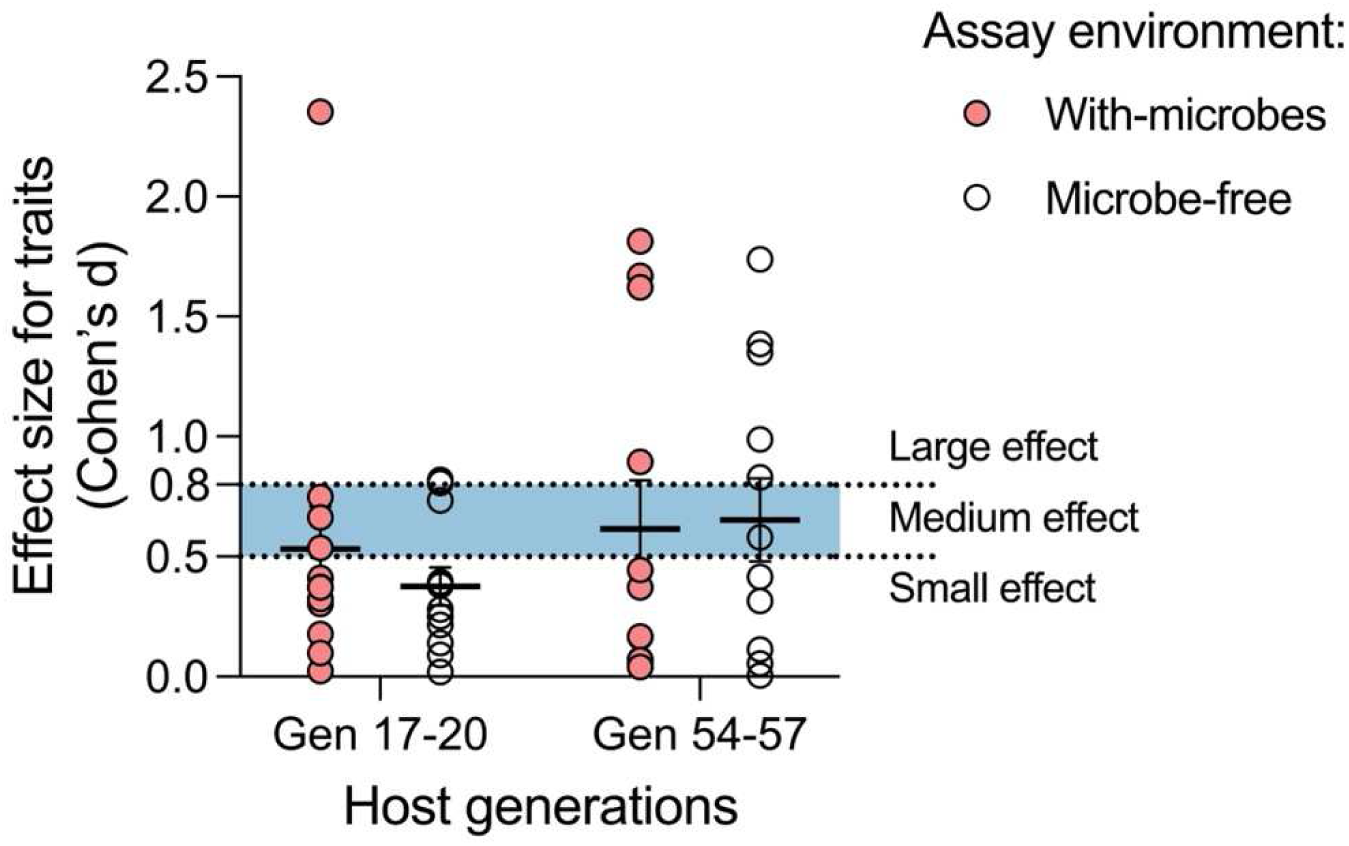
Changes in effect sizes (Cohen’s d) over all traits with time. Means are indicated by black horizontal lines. The error bars represent SEM. Coral circles indicate with-microbes environment and empty circles indicate the microbe-free environment.

The effect sizes (using Cohen’s d) for the phenotypic comparison between MB-MBLs in generations 17-20 were mostly small, with some effects being medium or large (Figure 6-left). In generations 54-57, the distribution of effect sizes shifted in the direction of large effects (Figure 6-right). This pattern is seen in both assay environments: with-microbes (circles with a coral fill) and microbe-free (circles without any fill). This suggests that the MBLs are still diverging from MBs even after 54-57 generations of experimental evolution.

### RNA-Seq results – Gen 54-57

After investigating the effects of long-term microbiome absence using a broad range of phenotypic assays, we used RNA-Seq to determine if specific genes are expressed differentially in MBL_1_ vs. MB_1_ as a direct consequence of their different evolutionary history.

In the microbe-free regime, where both selected and control populations were assayed without their microbiome, very few transcripts showed differential expression when analyzed using DESeq2 (Supplementary Text S7.1).

In the with-microbe regime, anti-microbial peptide (AMP) expression was up-regulated in MBL**_1_** compared to their control MB**_1_** (Figure 7). The complete list of up-regulated genes in MBL**_1_** is given in Supplementary Table S7. We also find that a cluster of heat shock proteins (HSPs) and several other genes in MBL_1_ are down-regulated. The complete list of down-regulated genes in MBL**_1_** is given in Supplementary Table S8. The gene ontology analysis for differentially expressed genes is presented in Supplementary Text S7.3.

**Figure 7.**
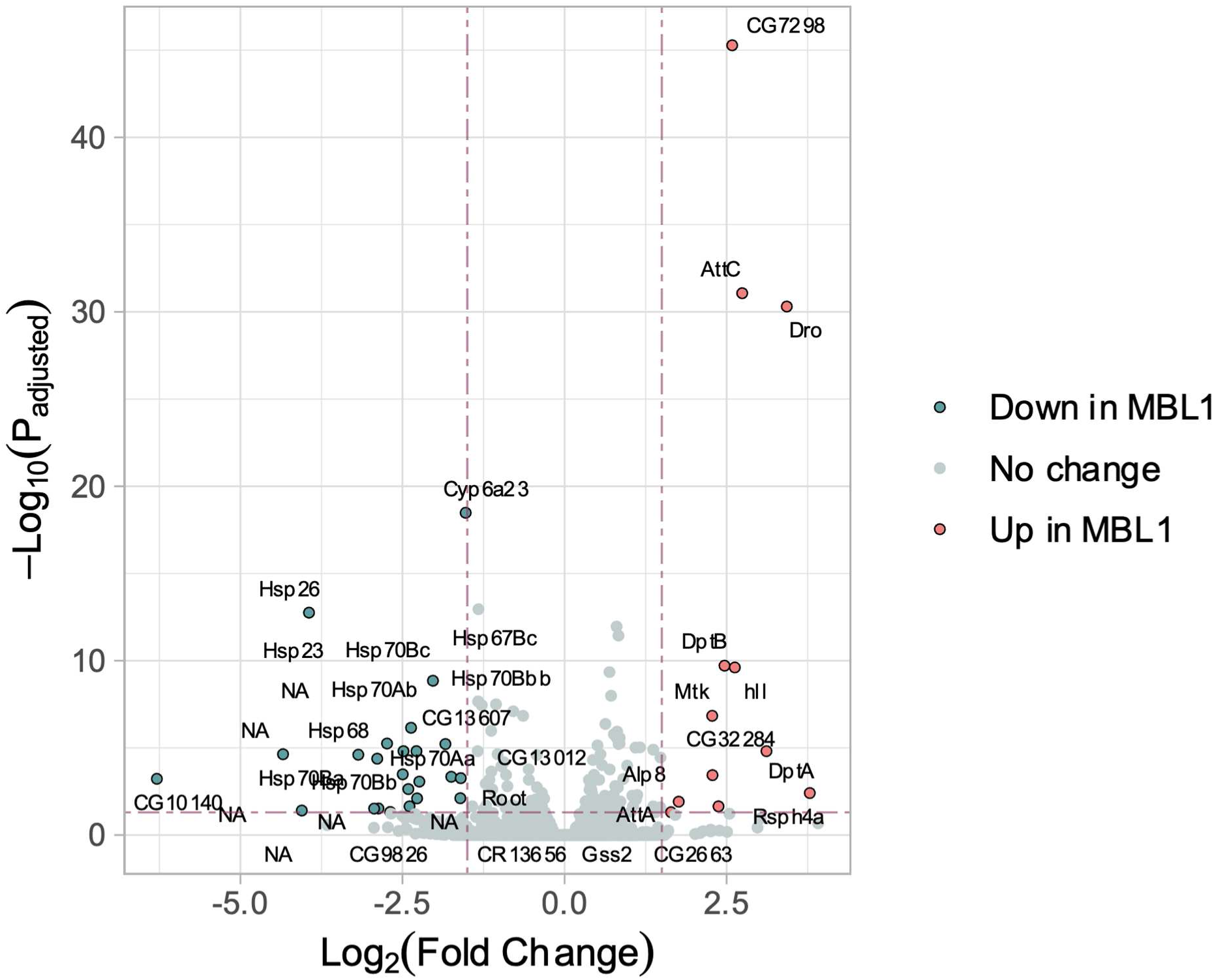
Differentially expressed genes in MBL_1_ vs MB_1_ in the with-microbes environment using DESeq2. The red dots represent the up-regulated genes, while blue dots represent down-regulated genes in MBL_1_. The protocol for the RNA-Seq is described in the method section.

Among all the differentially expressed genes, only those genes with Log_2_(Fold Change) > 1.5 and adjusted P-value < 0.05 are highlighted in Figure 7 and given in Supplementary Tables S7 and S8.

## Discussion

### MBL_1-4_ show few changes even after 54 generations without microbes

In a previous study using the ancestral outbred NDB_1-4_ flies (Malwade, 2024), it had been found that upon removal of microbiota, the female flies had lower fecundity, higher desiccation resistance, and lower locomotor activity, while for the males, locomotor activity increased. For both sexes, the other traits, such as body weight, egg-to-adult survival, mating latency, and mating duration, remained unchanged.

This led us to infer that microbiota plays an important role in the realization of several traits in *Drosophila melanogaster*, an observation that corroborates the results of numerous studies from the literature (Broderick & Lemaitre, 2012; Douglas, 2018b; Erkosar et al., 2013; Lesperance & Broderick, 2020; Ludington & Ja, 2020). This led to the hypothesis that the removal of microbiota is likely to be extremely stressful for the flies, as many traits would be affected at once. Hence, we expected the microbiota-less environment to exert very substantial selection pressure on the MBL_1-4_ flies, potentially leading to changes in multiple life-history traits, possibly including some trade-offs, within relatively few generations. However, the results of this study indicate otherwise.

In the first set of phenotypic assays done between host generations 17-20 (Supplementary Text S6), we saw that the mating time of MBLs was shorter than the MBs (Supplementary Figure S13D, with-microbes environment), and female body weight was slightly elevated for MBLs (Supplementary Figure S12A), microbe-free environment). Apart from these two changes, we did not see any evidence of evolutionary divergence between MB_1-4_-MBL_1-4_ in either of the two assay environments. This lack of divergence could be potentially attributed to the relatively short duration of the selection regime at that point.

In the second set of phenotypic assays done between host generations 54-57 (Figures 2 to 4), we saw that two traits showed significant change. In the microbe-free environment, we found that MBL_1-4_ had better egg-to-adult survival than MB_1-4_ (Figure 4A). In the with-microbes environment, MBL_1-4_ males had lower desiccation resistance (Figure 2C). It is possible that this lower desiccation resistance might be a cost incurred by MBLs in this environment.

When we looked at the effect sizes over all the traits assessed in Gen 17-20 vs. Gen 54-57 (Figure 6), we saw that the effect size distribution shifted from mostly “small and medium” effects to “small, medium, and large” effects. This suggests that even after 54 generations, the selected flies might still be diverging from the control flies.

We assayed the life-history traits both in the presence and the absence of microbes. Thus, any improvement in the fitness of the MBLs (compared to the MBs) in the microbe-free environment can be construed as the effect of adaptation in the selection environment. On the other hand, any reduction in the fitness of the MBLs (again compared to the MBs) in either assay environment would signify a cost of adaptation under microbe-free conditions. Our results showed very few adaptations or costs for the MBLs.

There are several potential ways, not all mutually exclusive, to interpret these observations. One possibility is that the selection pressure on the MBLs was so strong that they responded extremely fast, and therefore, by the time we assayed them first (i.e., by 17-20 generations), they had already responded maximally to the selection pressure and were back to the MB levels across all traits. However, if this were to be the case, in the microbe-free assay environments, we would have expected the MBLs to have generally greater fitness than the MBs. This is because the microbe-free assay environment is the “selection” environment for the MBLs but a novel environment for the MBs. Since the MBLs did not do better than the MBs in this assay environment, we conclude that the former had not adapted too well in the selection environment.

Another possibility is that even though the selection pressure is strong, our fly populations did not carry sufficient genetic variability to respond to the selection pressure. This might have been a valid concern if, like many studies on *Drosophila*-microbiota systems (e.g., (Ridley et al., 2013; Shin et al., 2011)), the experiments were performed on highly inbred or iso-female strains (e.g., *Canton-S, Oregon-R,* and *w^1118^*). However, our fly populations are outbred, are vigorously active in the cage, and have high fecundity and egg-to-adult survivorship (personal observations). Therefore, we have no reason to believe these populations lacked genetic variation. More importantly, the fact that egg-to-adult survivorship did change over time indicates that the populations were indeed evolving, which would not have been possible in the absence of genetic variation.

A third possibility is that while some divergence has indeed happened, they have not yet been picked up by our assays. For example, some previous studies have shown that the role of microbiota in affecting host physiology becomes prominent only in nutritionally unbalanced diets (Shin et al., 2011; Storelli et al., 2011). It is entirely possible that we will see more differences between the MBs and the MBLs if we assay them on a different diet. However, previous observations on these populations raised on standard food medium have shown that removing the microbiota leads to changes in multiple traits, even within a single generation (Malwade, 2024). Therefore, it is unlikely that the lack of differences between MBs and MBLs is due to the medium on which they were assayed or selected.

This brings us to the fourth possibility that although the removal of microbiota can lead to large-scale changes in a number of life-history and behavioral traits in a single generation, this does not translate into strong selection pressure on an evolutionary timescale. Our results would make sense when we consider the possibility that if the selection pressure on the host to evolve in the absence of microbes is low, then the evolutionary response can be slow and mild, as we observed in this study. We suggest that the results of our long-term experimental evolution are consistent with the so-called “evolutionary addiction hypothesis” (Douglas, 2018a; Hammer, 2024; Moran et al., 2019). This hypothesis suggests that when the microbiome is perturbed or culled entirely, hosts experience “withdrawal symptoms” as the host and microbes have lived together for an extended period (Hammer, 2024). Stated differently, although the host can function in the absence of the microbes, there is an inertia borne out of very long coexistence that temporarily affects the host. Thus, when we remove the microbiome for the first time, the host enters an altered state of physiology as it has already evolved some dependence on the previously ever-present microbes. Nevertheless, the host still has the capacity to perform its function without the microbiome, and within a few generations of microbe-free existence, it adjusts to the absence without any major phenotypic changes. This idea of evolutionary addiction might offer one potential explanation for the apparently low/weak selection pressure in our experimental evolution regime, and why we have not seen major evolutionary changes in our experimental evolution so far.

We speculate that the idea of evolutionary addiction is also consistent with the increased robustness seen in the MBLs (Fig 5). If growing in the absence of the microbes were simply helping the MBLs to come out of their “evolutionary addiction,” then one would expect that the microbiota would play a relatively lesser role in shaping their traits (relative to the MBs). In other words, the MBLs are expected to become more independent of their microbiota, increasing their robustness across the with- and without-microbes environments (by decreasing the sensitivity to microbiome presence/absence). Both proxies of robustness, Cohen’s d (Fig 5A) and percentage change (Fig 5B), suggested that this is the case. However, the effect size in both cases was small, which suggests that the MBLs still have some distance to go to “kick their addiction,” to function fully independent of the microbiome, and the phenomenon is perhaps incipient at this stage.

We note here that our conceptualization of robustness is close to phenotypic plasticity. However, phenotypic plasticity is typically defined on genotypes (Agrawal, 2001; Sommer, 2020), whereas our traits were measured at the population level, which likely consisted of multiple genotypes. That is why we refrained from calling our MBL populations less plastic. However, our observations on robustness do raise the possibility that the plasticity of the MBL populations might have been altered due to selection. This conjecture needs to be examined in detail in future studies.

### MBL_1_ and MB_1_ have at least three sets of differentially expressed gene clusters

To see if MBLs have started showing differences in gene expression compared to MBs, we performed an RNA-seq on one group of populations, namely MBL_1_ and MB_1_. We here note that these findings, on only one of the four evolutionary replicates, are preliminary.

#### A cluster of heat shock proteins (HSPs) is downregulated in MBL_1_

Heat shock proteins are the proteins that are involved when the organism experiences stress (Feder & Hofmann, 1999). Even though, as their name suggests, they play an important role primarily in the heat shock response (Lindquist, 1986), some of these proteins can be expressed in response to various other stresses such as cold, crowding, anoxia, desiccation, etc., where these proteins can act as chaperones (King & MacRae, 2015). We find that HSPs (*HSP23, HSP26, HSP67, and HSP68*) and HSP subunits (*HSP67 and HSP70*) are down-regulated in MBL_1_ flies that were maintained without the microbiome for multiple generations.

#### RNA-Seq results on MBL_1_ might be consistent with the desiccation resistance trends over all four replicate MBLs

Lower expression of HSPs in MBL_1_ suggests a potentially muted stress response of the MBLs due to the lower availability of HSPs to counter the stress. This correlation of stress resistance with the expression of HSPs is well-documented (Feder & Hofmann, 1999).

When we looked at the desiccation resistance of MBL_1-4_ vs. MB_1-4_ in the with-microbes regime, we found that MBL_1-4_ had a trend of lower desiccation resistance than MB_1-4_ for males (Figure 2C) but not for females (Figure 3C). Both these effects were large in size. While we have not established a causal link in this trend of lower desiccation resistance, this observation could be a potential link between the lower expression of HSPs and lower desiccation resistance.

Looking at this desiccation-HSP expression correlation, one can speculate that the thermal stress tolerance of MBL_1-4_ will be lower than that of MB_1-4_ due to lower expression and, hence, lower levels of HSPs. But that need not be necessarily true. A study of land snails has shown that desiccation-sensitive species maintain more HSPs than desiccation-resistant species (Mizrahi et al., 2010). If the same logic holds for thermal stress tolerance in *Drosophila*, then the lower expression of HSP might hint at a better ability of MBLs to resist heat stress. Thus, from this data, we cannot predict the ability of the MBLs to withstand thermal stress.

#### Antimicrobial peptides (AMPs) are upregulated in MBL_1_

Antimicrobial peptides (AMPs) are small cationic molecules that interact with bacterial membranes to damage and kill the bacteria (Brogden, 2005). They are widespread across the tree of life, with animals and plants harboring various types of AMPs to deal with pathogens (Mookherjee et al., 2020). In *Drosophila melanogaster*, there are seven types of AMPs whose functions are known, and a detailed discussion on them is available elsewhere (Hanson & Lemaitre, 2020; Lemaitre & Hoffmann, 2007). In addition to their classical role of defense against pathogens, these AMPs are also shown to control the host microbiome (Franzenburg et al., 2013; Maritan et al., 2024; Marra et al., 2021; Mergaert, 2018).

A study that looked at the gene expression changes in the *Drosophila* gut using transcriptomics when the host is colonized with microbes vs. when it is kept without microbes found that the presence of a microbiome leads to upregulated expression of several immune response genes, including AMPs such as *Attacin*, *Drosomycin* and stress response genes such as *HSP23*, *HSP67Bb*, and *HSP70Bc* (Broderick et al., 2014). While their observation of the up-regulation of the immune response matches our results in MBL_1_, the HSP response is down-regulated in MBL_1_, in contrast to the observation by Broderick et al., which was done over the host’s lifetime (Broderick et al., 2014). The up-regulation of the immune system upon microbiome colonization is also reported in other subsequent single-generational studies that are done over the host’s lifetime (Chandler et al., 2022; Elya et al., 2016).

The above single-generation studies on *Drosophila* indicate that the presence of a microbiome leads to AMP up-regulation compared to when the microbiome is removed. In our study, as both MB and MBL received the same native microbiome from a common pool (shown in Supplementary Text S5.1-S5.2), their different evolutionary history should be the driving force in the up-regulation of AMPs, not the differential presence of microbiome.

#### A hygiene hypothesis-like phenomenon can potentially explain the overactive humoral immune response in MBLs

MBLs have been kept away from any microbes for more than 50 generations. Several studies have shown that microbes are indispensable for the proper calibration and modulation of the immune system, especially early in life (Abt et al., 2012; Donald & Finlay, 2023; Gensollen et al., 2016; Henrick et al., 2021; Hoang & King, 2022; Knoop et al., 2017; Mazmanian et al., 2005; Metcalf et al., 2022; Selosse et al., 2014). This calibration includes proper sorting of self vs. non-self (W.-J. Lee, 2008; Lhocine et al., 2008). There is a possibility that MBLs might have lost the delicate balance that maintains this calibration and, as a result, are over-expressing AMPs compared to MBs to actively regulate the microbiome. This possibility can be validated in future experiments.

In humans, this phenomenon is often discussed in the context of the hygiene hypothesis (or a refined version of the same, called the old friend hypothesis) (Bach, 2018; Blackwell, 2022; Wills-Karp et al., 2001). This hypothesis states that exposure to certain microorganisms, particularly in early life, can reduce the frequency of atopic diseases and immune-mediated disorders in organisms (Stiemsma et al., 2015). It is possible that the observed increased levels of AMPs signify immune overactivation via immune miscalibration over many generations. It is already known that *Drosophila* can suffer from autoimmune disorders (Mortimer et al., 2021). Therefore, if our conjecture about the hygiene hypothesis is true, then one might predict that the MBLs would suffer from higher instances of autoimmune disorders, which we would like to investigate in detail in a future study using all four evolutionary replicates.

Apart from the HSPs and AMPs, another smaller set of chitin-binding genes is also differentially expressed in MBL_1_. We have discussed this in Supplementary Text S8.

We have discussed multiple speculative insights in the latter part of the discussion, even though they are not substantially supported by our current data, as experimental microbiome manipulation studies on host-microbiome long-term interactions are rare and demanding, and at least some of these insights that we discussed might help generate further falsifiable hypotheses in *Drosophila* and in other hosts.

## Conclusion

Our study involved *Drosophila melanogaster,* where the short-term removal of microbiota is known to have major effects across multiple traits (Gnainsky et al., 2021; Jia et al., 2021; Schretter et al., 2018; Shin et al., 2011; Storelli et al., 2011; Suyama et al., 2023). Therefore, we expected that the removal of the microbiota for an extended period would force the populations to evolve along multiple trait axes, potentially with associated life-history costs. Contrary to our expectations, even after 54 generations, there were only modest evolutionary changes in the host. This might be due to weak evolutionary pressure on the host and may be situated in the framework of the “evolutionary addiction hypothesis.” This might also mean that, for the outbred *Drosophila* system, the “host + microbe” unit (i.e., the holobiont) might represent a somewhat weak integration. Based on our results, we cannot claim that the cost of evolution in the absence of the microbiome will be negligible in other systems. However, our results suggest that if the host pays large costs of microbiota removal across multiple traits in a single generation, then that does not necessarily mean that the host is under strong selection pressure and will evolve to counter this pressure in the long run. In fact, very little can be predicted about the long-term effects of evolution under a no-microbe condition based on insights from the existing single-generation studies. Previous theoretical (Henry et al., 2021; Vliet & Doebeli, 2019) and empirical (reviewed in (Rosenberg, 2021)) studies have suggested that microbiota can affect the evolutionary trajectories of their hosts. Our study suggests a more nuanced picture. Taken together, our findings indicate that the role of microbiota in determining long-term evolutionary trajectories deserves further critical examination.

## Author contributions

AM and SD formulated the study. AM, AK, BV, CG, SS, and VK conducted the assays. AM and SD did the data analysis. AM and SD wrote the manuscript with inputs from the other authors.

## Acknowledgments

We acknowledge the help of Krishna Chaitanya Sattaru, Navanath Kadam, Saee Patil, Shreya Mishra, Sohum Ranade, Viraj Gaonkar, and Yashvita Subramanian in data collection. We thank Amitabh Joshi, Deepak Barua, and Yashraj Chavhan for helpful discussions and suggestions related to this project.

Data availability:

All the data relevant to this study will be made publicly available in the Dryad Digital Repository upon acceptance.

## Supplementary information

**Figure S1.**
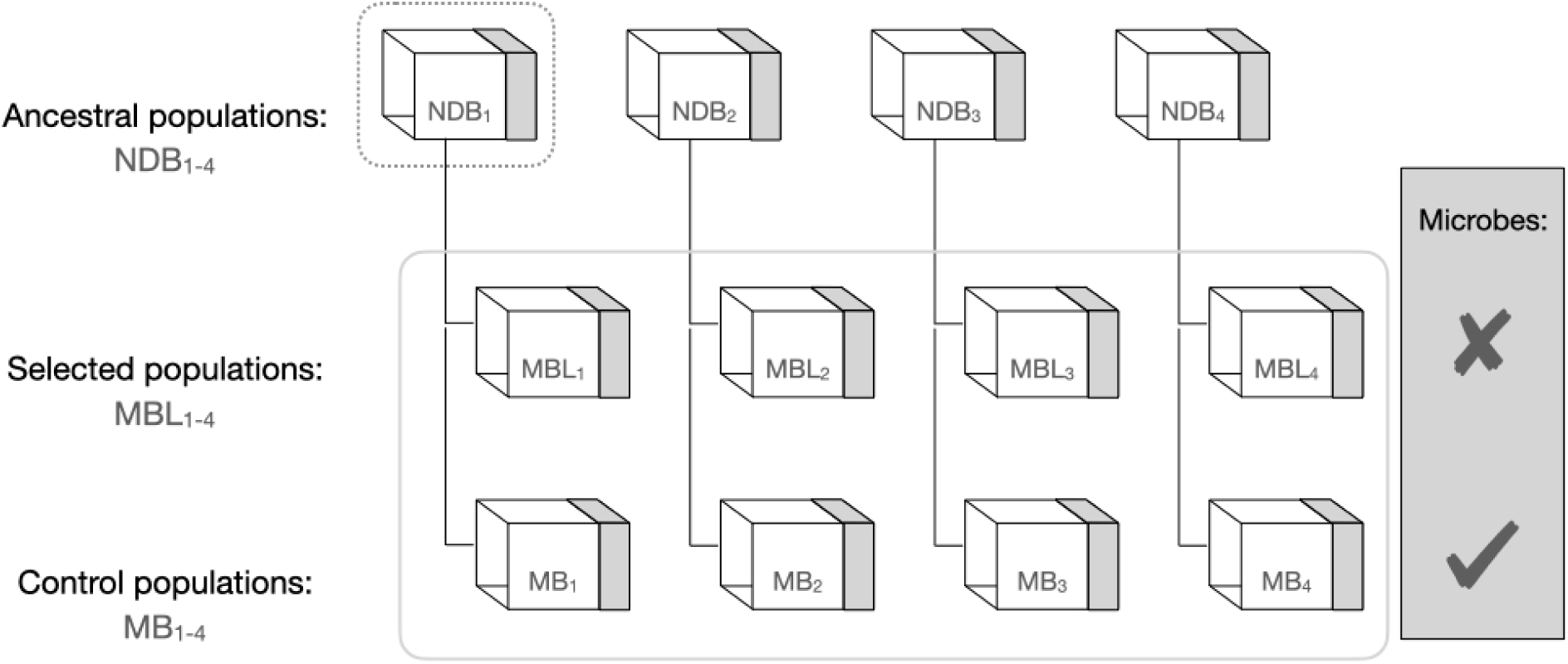
Schematic showing the design of the study.

**Figure S2.**
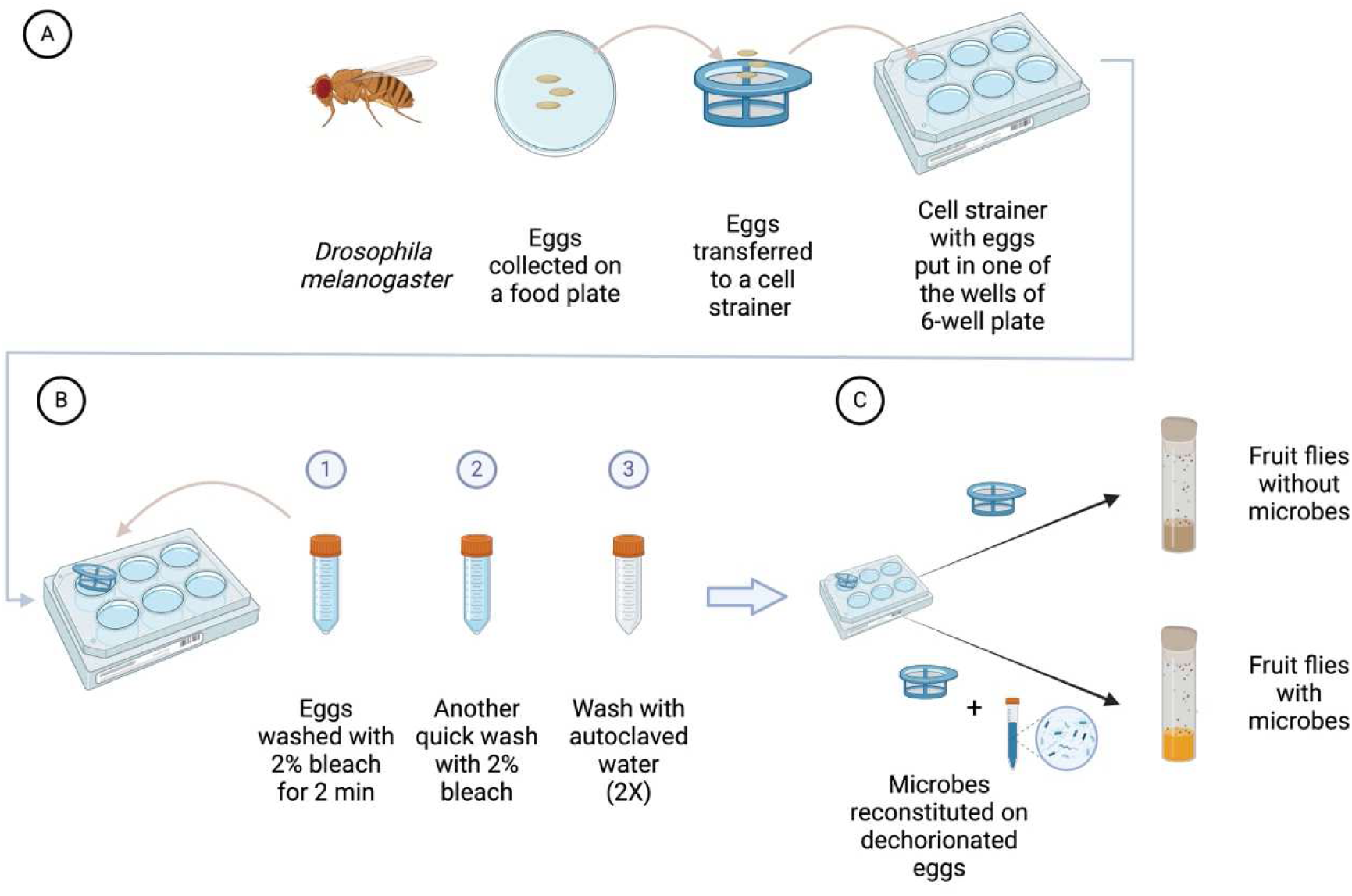
Details of the egg dechorionation protocol for making MBL1-4 and MB1-4 populations. Created in BioRender.com.

**Figure S3.**
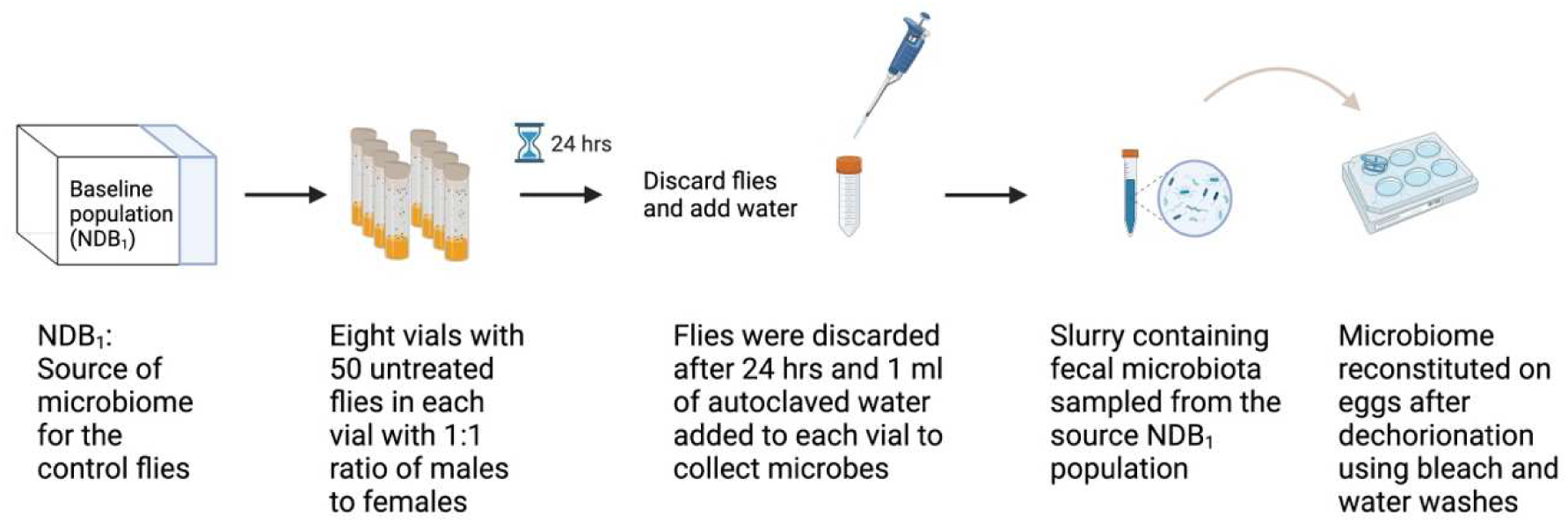
Schematic of the detailed protocol of microbiome reconstitution from NDB1. Created in BioRender.com.

## Supplementary information

### Text S1 Details of the baseline populations and amplicon sequencing

#### S1.1 *Drosophila* NDB stocks and rearing

Wolbachia-free *Drosophila melanogaster* used in the single-generation study were derived from four large, outbred laboratory populations named New Dey Baselines (NDB1-4). These fly lines were maintained on the standard banana-jaggery food with a 14-day discrete generation cycle at 25°C under constant light conditions with a breeding population of ∼2400 individuals. NDBs trace their ancestry to the DB populations (see Supplementary Text S1.3 for the details), which in turn trace back to the wild-caught IV lines (Ives, 1970; Rose, 1984) via the JB1-4 populations (Sheeba et al., 1998).

Pairs of MB-MBL populations that have the same subscript are ancestry-matched, and are referred to as “block” in this document (e.g., MB1 and MBL1 are together called block 1).

#### S1.2 Maintenance regime of baseline *Drosophila* populations (NDB1-4)

In the lab, the adults of the four populations (NDB1-4) are maintained in plexiglass cages (25 cm x 21 cm x 16 cm) at a large population size of ∼2400 per population. On day 12 after egg collection, the fly population is provided with a moist yeast paste on top of the banana-jaggery food to boost egg output. This yeast paste also contains a few drops of glacial acetic acid to attract the flies to the paste. At the end of day 13, a food plate, with vertical edges of food exposed for oviposition, is provided for 12-16 hrs. On day 14, deposited eggs are collected by cutting thin food strips from the exposed surface, under a binocular stereo microscope. The size of the strips is adjusted under the microscope using a clean scalpel such that each strip contains about 400 eggs. These strips are then transferred to plastic and translucent *Drosophila* milk bottles containing ∼50-60 ml of banana-jaggery food at the bottom. This marks the new generation’s 0th day after egg collection. Adult flies arising from six such bottles are transferred to fresh plexiglass cages on the 12th day (post egg collection). This gives rise to a new generation with a population size of ∼2400 individuals, which do not coexist with the adults of their parental generation at any stage.

#### S1.3 *Drosophila* DB stocks and rearing

NDB1-4 were derived from their most recent ancestors in our lab, Dey Baselines (DB1-4) (Sah et al., 2013). These four DBs were maintained in our lab for 220 generations and were used as baselines for this period. We pooled an equal proportion of eggs from each DB population and created four independent NDBs. We maintained NDBs separately for ∼10 generations without any assays for them to stabilize after this intermixing. The maintenance of DBs was similar to NDBs except for two differences: (a) DBs were maintained in *Drosophila* 8-dram vials at an egg density of ∼ 60 per vial with ∼ 6 ml of food as opposed to NDBs that are reared in milk bottles (b) DBs were yeasted on 18th day, provided with a cut plate on 20th day, and their eggs were collected for next generation on day 21. Thus, DBs followed a 21-day discrete generation cycle as opposed to a 14-day discrete generation cycle followed by NDBs.

#### S1.4 Population-level 16S rRNA amplicon sequencing

To know the relative composition of bacteria in *Drosophila* populations, DNA was extracted as described in Supplementary Text S3.2.2. DNA was run on 1% agarose gel to see the extraction quality. After the integrity check on agarose gel, DNA was sent for 16S rRNA V3-V4 region amplicon sequencing to a commercial vendor (1st BASE Laboratories, Malaysia). Sample quality was again confirmed by the vendor using agarose gel, spectrophotometer, and fluorometric method. All samples passed DNA quality check and were subjected to amplicon PCR followed by a quality check. These samples were then processed to prepare amplicon libraries using 2-step PCR according to Illumina’s 16S metagenomic library preparation guidelines (Link to the guidelines - https://sapac.support.illumina.com/downloads/16s_metagenomic_sequencing_library_preparation.html). These libraries were processed using Illumina’s MiSeq platform using 2x300bp chemistry.

#### S1.5 Data analysis of amplicon sequencing

Sequence adapters and low-quality reads were removed from the paired-end reads before the first 2,00,000 raw reads were extracted using BBTools (https://sourceforge.net/projects/bbmap/). The forward and reverse reads were merged using QIIME (Bolyen et al., 2019). DADA2 pipeline (Callahan et al., 2016) was used to remove low-quality regions and chimeric errors. The resulting data was in the form of amplicon sequence variants (ASVs). The taxonomic classification was done using scikit-learn (Pedregosa et al., 2011) and the Naive Bayes classifier against the SILVA database (release 132). This part of the analysis was performed by the commercial vendor (1st BASE Laboratories, Malaysia).

#### S1.6 Details of RNA-seq

##### RNA isolation

Total RNA was extracted using miRNeasy Mini kit (Qiagen, Catalog No. 1038703) following the manufacturer’s protocol. About 30 females were homogenized and lysed in Trizol reagent, followed by chloroform extraction. The aqueous phase was mixed with 1.5x chilled ethanol, and the mix was loaded on the columns provided in the kit. RNA was eluted in 30 µl RNase-free water. The RNA samples were quantitated on a Qubit (Thermo Fisher Scientific) fluorometer. Appropriate dilutions were loaded on a high-sensitivity RNA screen tape to determine the RNA integrity number (RIN).

##### Library preparation and sequencing

1.5 µg of RNA per sample was depleted of fly rRNA using QIAseq FastSelect - rRNA Fly Kit (Qiagen, Catalog No. 333262). The NEBNext Ultra II directional RNA library prep kit (Illumina, Catalog No. E7760L) was used to construct double-stranded cDNA libraries from the rRNA-depleted RNA. The cleaned libraries were quantitated on a Qubit fluorometer, and appropriate dilutions were loaded on a high-sensitivity D1000 screen tape to determine the library size.

These libraries were sequenced on the Illumina NGS platform with 2x150bp chemistry to generate 10 million paired-end reads (3 Gb data per sample).

##### RNA-seq analysis

The quality of the reads was assessed using *FastQC v0.11.9* (Andrews, 2010). These reads were passed through *fastp* to remove adaptors and low-quality bases. The filtered reads were mapped against the *Drosophila melanogaster* BDGP6.46 top-level chromosome (GCA_000001215.4) retrieved from ENSEMBL using *HISAT2* (Kim et al., 2019). The program *featureCounts* (mode = fragment) was used to count mapped reads. The count table from *featureCounts* was given as input to *DESeq2* (Love et al., 2014) to find differentially expressed genes in the populations.

Transcripts with counts less than ten were removed from all samples. The false discovery rate (FDR) cut-off was at 0.05, and the log fold change (LFC) threshold was at 1.5.

### Text S2 - Note on the use of F1 flies for the assays

In our flies, we removed microbes via dechorionation at the egg stage. Even if we have taken care of the confounding effect of bleach by using proper control, we wanted to keep the effect of the bleach on flies to a bare minimum (ideally none). To that end, we employed an additional measure of using F1 flies for the assays, as explained below (Supplementary Figure S4).

In this approach, to generate flies without the microbes, we followed a protocol employed by several other studies where F1 flies (progeny of the flies that experienced the bleach at the egg stage) were used in the assays (H.-Y. Lee et al., 2019; Pais et al., 2018; Selkrig et al., 2018). This ensured that the F1 assay flies of both types, microbiome-reconstituted (M+) and microbe-free (M-), did not experience bleach in their lifetime (their respective parent population did). To generate F1 M- assay flies, F1 eggs were collected from the flies without microbes (i.e., F0 M- flies, Supplementary Figure S4) and were raised in aseptic conditions, ensuring the absence of microbes. To generate control F1 M+ assay flies, F1 eggs were collected without any manipulation from the flies whose microbiomes were reconstituted (i.e., F0 M+ flies, Supplementary Figure S4).

**Figure S4.**
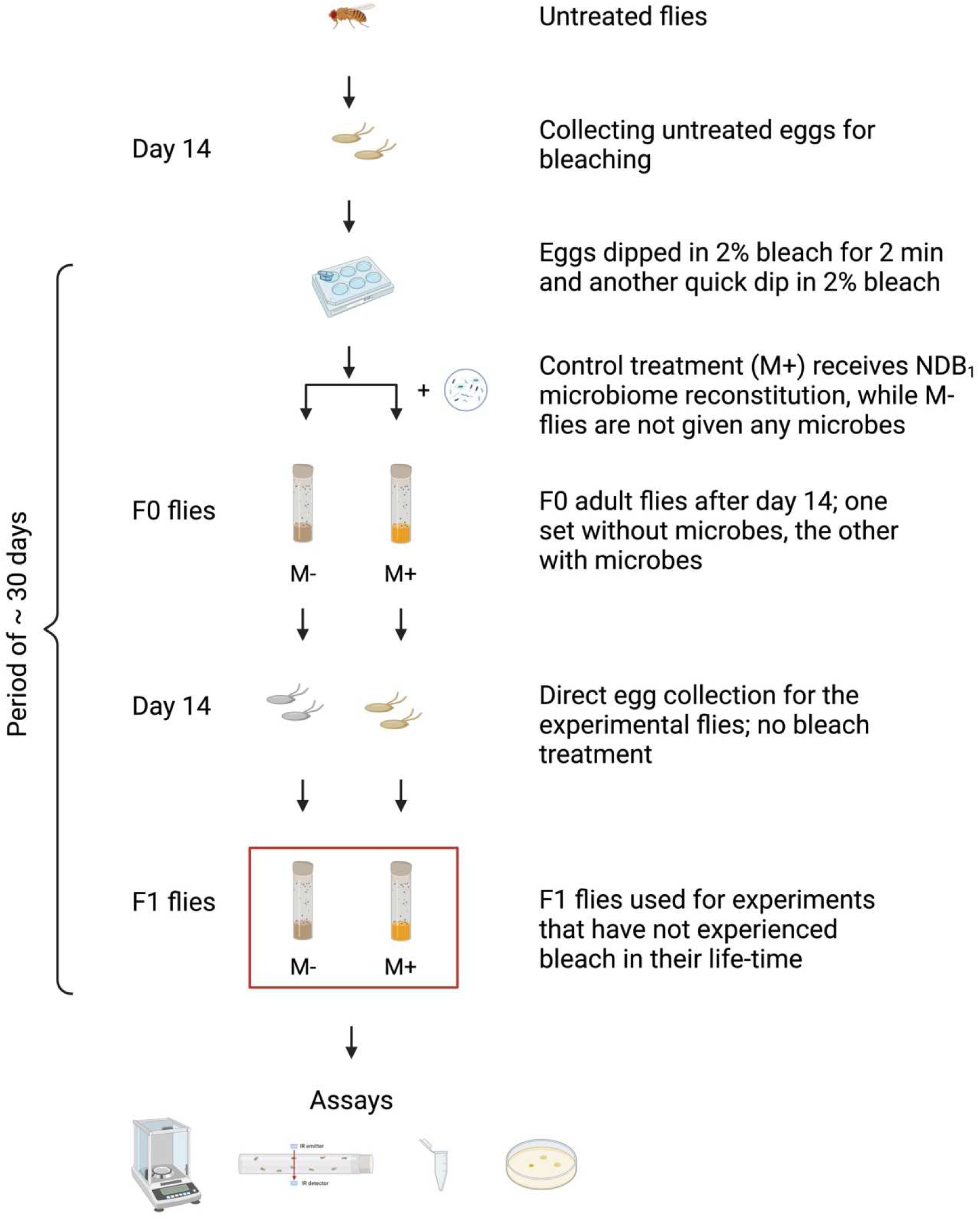
Schematic of the full experimental design to study microbiomes’ contribution to host phenotypes. To create two types of flies (with the minimal effect of bleach), flies with microbes (M+) and without them (M-), we combined two procedures. First, we chose microbiome-reconstituted flies as a control instead of using untreated flies. The F0 flies of both (M+ and M-) types were generated using bleach at the egg stage. This protocol is illustrated in detail in Figure S2. Second, as discussed in Supplementary Text S2, assays were conducted on progeny of these F0 flies (i.e., F1 flies) that have not experienced bleach in their lifetime to avoid any recency effect of bleach. The eggs collected for F1 experimental flies either had the microbes (M+) or lacked the microbes (M-) depending on the state of their parent’s microbiome. Created in BioRender.com.

### Text S3.1 - Assessing the microbiome status of MBs and MBLs during experimental evolution

We plated adult flies from MBL1-4 every generation to see if bacteria are indeed absent in these populations (data not shown here). Except for generations and blocks specified in Supplementary Table S1, in all other generations, MBLs did not show the presence of a microbiome. We sometimes did observe ∼1-2 colonies (per 10 flies plated) in some replicates.

**Table S1.** Generations in which MBLs showed transient contamination.

| Generation | Affected MBL blocks |
| --- | --- |
| 37 | Block 1 and 3 |
| 38 | Block 1 |
| 39 | Block 1 |

MB1-4 were plated every 2-3 generations and CFUs were counted to make sure they received the reconstituted microbiome (data not shown here).

To confirm that our protocol indeed removed both culturable and unculturable microbes, we performed PCR with 16S rRNA universal primers (Supplementary Text S3.2.2) on DNA extracted from adult MB1-4 and MBL1-4 females (day 15 after egg collection) in generation 40. In Supplementary Figure S5, we see that MB1-4 show the presence of the reconstituted microbiome, and the microbiome is absent from MBL1-4. This also confirms that, by generation 40, the contamination that we saw in MBLs in previous generations was completely gone.

**Figure S5.**
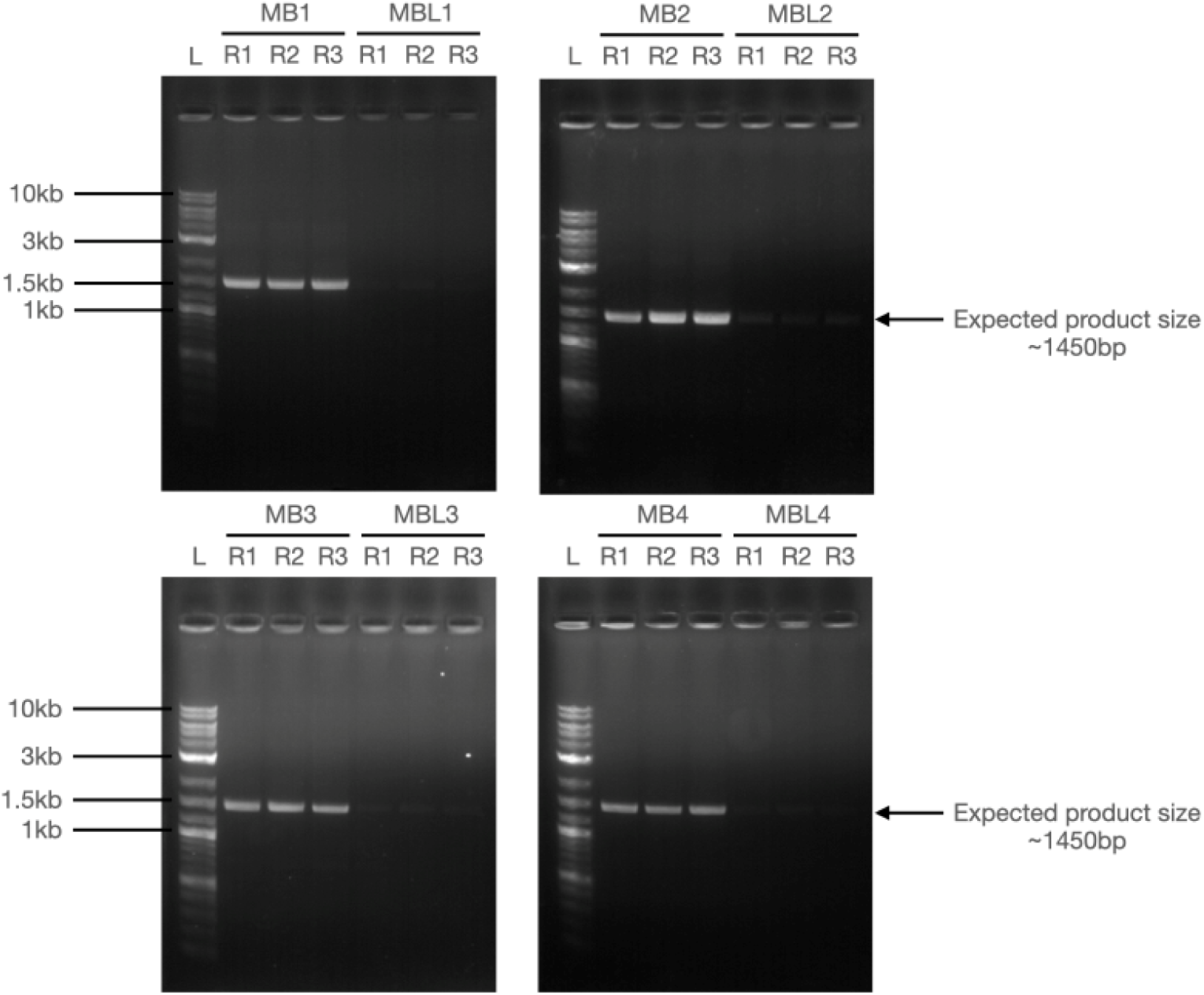
16S rRNA PCR to check the status of microbiome in MB1-4 and MBL1-4.

### Text S3.2 Fly-associated bacterial load analysis S3.2.1. Culture-dependent method

Adult flies (day 12 after egg collection, unless stated otherwise) were sorted by sex under mild CO2 anesthesia. Five male and five female flies under anesthesia were surface sterilized with 70% ethanol in 1 ml microcentrifuge tubes. These flies were then homogenized in 200 µl of autoclaved dH2O and plated on the nutrient agar (HiMedia) or the de Man, Rogosa, and Sharpe (MRS) medium (HiMedia) and incubated for 72 hrs and 48 hrs, respectively, at 25^0^C under aerobic conditions. Dilutions with 10-100 colonies were counted, and the counts were multiplied by the overall dilution factor to estimate colony-forming units (CFUs) per fly.

#### S3.2.2. Culture-independent method

To test the bacterial load using PCR, total DNA was extracted from eight to ten adult flies using the QIAamp DNA Mini kit (Qiagen, Catalogue No. 51304) as per the manufacturer’s instructions. We modified three steps of the manufacturer’s protocol in line with the protocol from an earlier study (Jehrke et al., 2018). First, we used mechanical homogenization (with a homemade handheld motorized pestle mixer) to lyse the tissue in 1.5 ml Eppendorf tubes instead of just Proteinase-K treatment. Second, 20 µl of 20 mg/ml RNase A (HiMedia, Catalogue No. DS0003-5ML) was added to each sample after tissue lysis. Third, the sample was incubated with “Buffer AL” at 70^0^C for 30 min instead of 10 min. We avoided any vortex step after tissue lysis to minimize the shearing of the DNA. The quality of total DNA was checked using agarose gel electrophoresis. This DNA, composed of the fly and microbial genomes, was used as a template for PCR using 16S bacterial universal primers with a product size of about 1.5 kb. The details of the primers are given below in Supplementary Table S2. The primers were ordered from Sigma-Aldrich, India. The PCR reactions were set up using GoTaq® Flexi DNA Polymerase (Promega, Catalogue No. M829) with 40 ng gDNA input. The initial denaturation was at 94^0^C for 2 min, followed by 30 cycles of (denaturation at 94^0^C for 30s, annealing at Tm = 60^0^C for 1 min, extension at 72^0^C for 1 min), with final extension 72^0^C for 10 min. The PCR products were visualized on 1% agarose gel with 1kb plus DNA ladder (NEB, Catalogue No. N3200S) as the molecular weight marker.

**Table S2.**
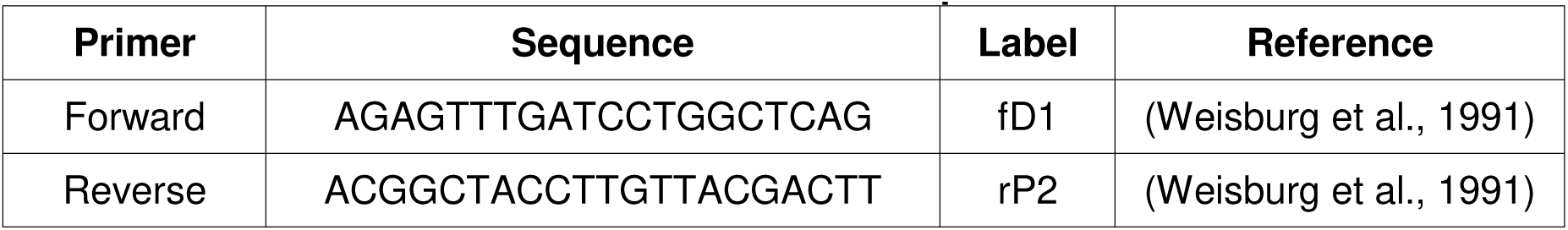
16S bacterial universal primers used for PCR.

### Text S4.1 – First phenotypic assessment of experimental evolution lines at host generations 17-20

After 17 generations of selection, we looked at various host traits to see if the experimental evolution without the microbes has led to any divergence in MBLs vis-à-vis the MBs. This assessment was done under two assay environments: (1) with-microbes and (2) microbe-free.

In this design, logistically, it was not possible to assay all four blocks of MB and MBL together. Hence, we assayed them in sets of two blocks, namely 1 - 2 and 3 - 4. Supplementary Table S3 gives details of the generations in which the assays took place. MB and MBL populations in each block were always assayed together in the same generation to know the effect of the selection procedure (e.g., MB1 compared with MBL1). For all assays done in this set, flies for separate environments (i.e., with-microbes and microbe-free) were generated in different host generations (Supplementary Table S3). As the two different environments were not assayed in the same host generation, we do not discuss the effect of the environment on these populations. We only discuss the effect of selection in this first set of assays.

**Table S3.**
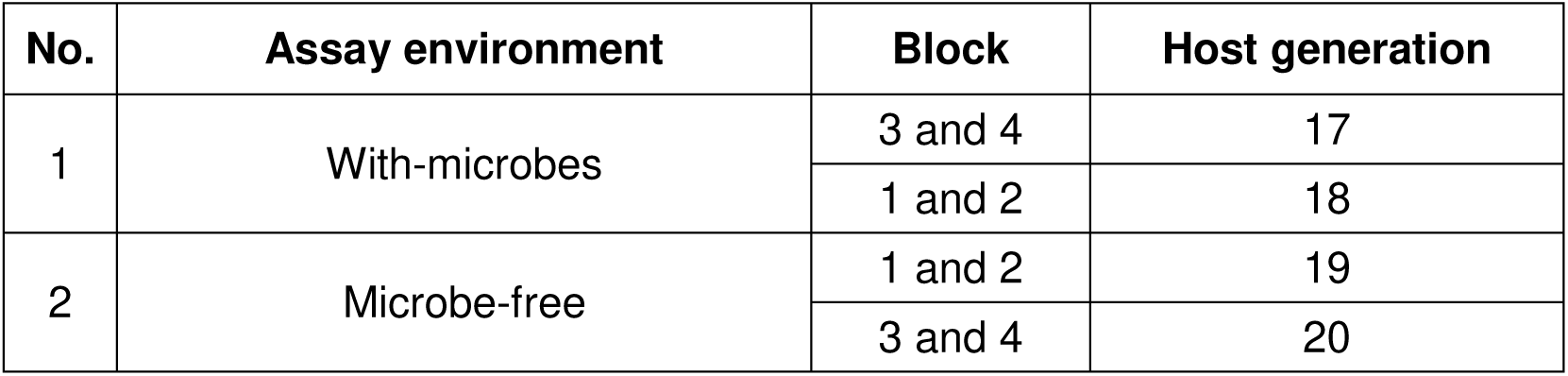
Details of the assay environments and host populations for Gen 17-20 assays.

The validation of the microbiome states of flies raised in with-microbes and microbe-free assay environments is reported in Supplementary section S5.1.

### Text S4.2 - Second phenotypic assessment of experimental evolution lines at host generations 54-57

In the first assessment of MBL1-4 vs. MB1-4 that happened in generations 17 to 20 (Supplementary Text S4.1), we noticed that the assay environments (with-microbes and microbes-free) had a greater effect in determining the trait-values in both sets of populations than the selection regime (Figures S11-S13). However, since the with-microbes and microbe-free flies were generated in separate host generations (see Supplementary Table S3), we could not compare the relative effects of assay environment vs. selection quantitatively with this design. Therefore, during the next set of assays during generation 54-57, we amended our experimental design by conducting the assays in the two environments simultaneously. This allowed us to directly compare the contributions of selection vs. assay environment. Supplementary Table S4 shows the details of our experimental protocol for the set of assays conducted during generations 54 to 57. This amendment meant, due to logistical reasons, we could only assess one block at a time in the same host generation (in contrast to the Gen 17-20 assays where two blocks were assessed in the same host generation).

This experimental design allows us to make two kinds of comparisons: (1) the effect of experimental evolution in a common-garden assay environment obtained by directly comparing MBs vs. MBLs (Main text, Figures 2 to 4) and (2) the effect of the presence vs. absence of microbes on both MB and MBL populations (Main text, Figure 5)

**Table S4.** Details of the assay environments and host populations for Gen 54-57 assays.

| No. | Common-garden environment | Block | Host generation |
| --- | --- | --- | --- |
| 1 | With-microbes | 1 | 54 |
|  |  | 2 | 55* |
|  |  | 3 | 56 |
|  |  | 4 | 57 |
| 2 | Microbe-free | 1 | 54 |
|  |  | 2 | 55* |
|  |  | 3 | 56 |
|  |  | 4 | 57 |

\*For block 2, body weight and female fecundity were assessed in Gen 59

### Text S5.1 - Validation of microbiome status in the flies raised in two different assay environments

#### S5.1.1 By plating fly homogenates

In the with-microbe environment, on both MRS and NA media, we see the presence of microbes (Supplementary Figure S6, Set-B and Set-C). Set-A is the autoclaved water that is used to homogenize the flies. The serial dilutions with ∼10-100 colonies were counted to get the CFUs/fly. While Supplementary Figure S6A shows a representative picture from block 1, CFUs/fly for all four blocks of MBs and MBLs are compared using paired t-test on block means in Figure S7.

In the microbe-free environment, both MB1 (Supplementary Figure S6B) and MBL1 (Supplementary Figure S6C) show the absence of the microbiome. While block 1 plates do not have any microbes, other blocks show a few colonies in some replicates (data not shown here).

**Figure S6.**
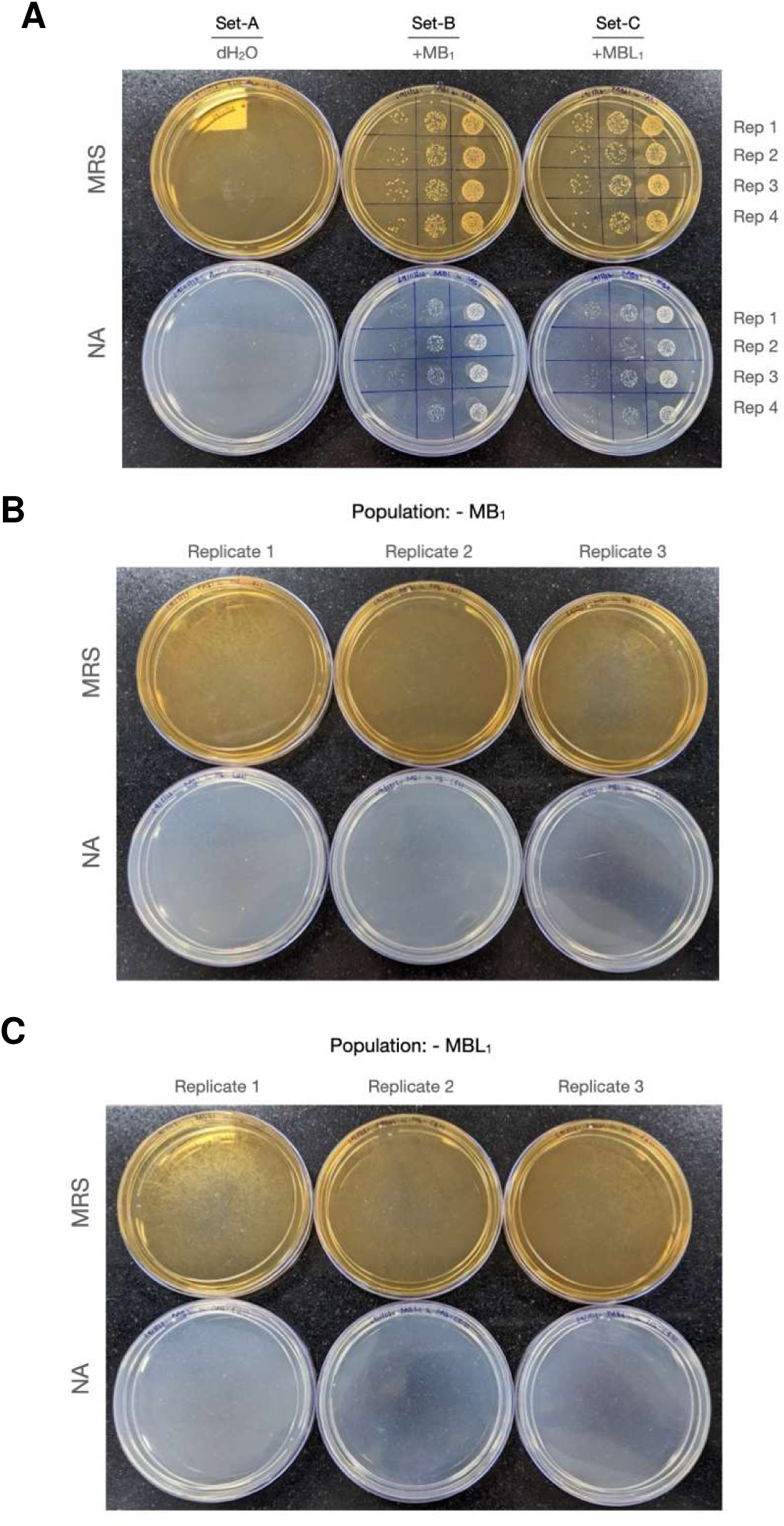
MRS and NA plates showing presence of microbes in with-microbe environment and their absence in microbe-free environment. (A) MB1 and MBL1 in with-microbe environment (n = 4). The spots show different serial dilutions used to calculate CFUs/fly. These CFU/fly for all the replicate populations are shown in Supplementary Figure S7. (B) MB1 microbe-free environment (n = 3). (C) MBL1 microbe-free environment (n = 3).

#### S5.1.2 MBs and MBLs received similar microbial loads in the with-microbe environment (Gen 54-57 assays)

**Figure S7.**
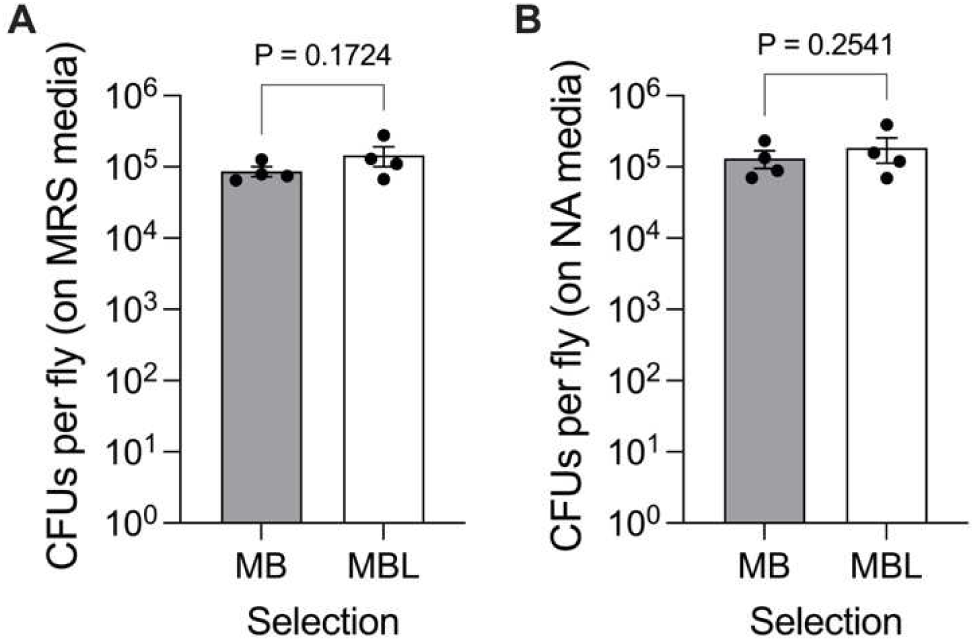
Bacterial load in the MB1-4-MBL1-4 flies (with-microbe environment) as determined by plating fly homogenates. CFUs/fly on (A) MRS-agar (B) NA-agar. The bacterial load in MBs vs. MBLs is compared using paired t-test on block means. Block means are indicated by black filled circles. The error bar represents SEM. We see that the process of microbiome reconstitution leads to MBs and MBLs carrying similar bacterial loads.

#### S5.1.3 PCR showing the presence of microbiome in the with-microbiome environment and depletion of the microbiome in microbe-free environment

To validate the status of the microbiome in both environments with a culture-independent method, we extracted DNA from the four types of populations (Main text, Figure 1B). We performed PCR with 16S rRNA universal primers (details in Supplementary Text S3.2). For PCR, a subset of flies used for the assay on day 12 were flash-frozen in liquid nitrogen and kept at -80^0^C for further use. For the with-microbe environment, 12 female flies from only MB1 and MBL1 (block 1) were used for the DNA extraction. For the microbe-free environment, total DNA was extracted from 12 female flies (3 flies per block, four such blocks). The PCR was done with 40ng of input DNA with 30 cycles.

In the with-microbe environment, MB1 and MBL1 showed the presence of microbes as expected (Supplementary Figure S8). In the microbe-free environment, while two replicates have no band, one replicate each from pooled MB1-4 and MBL1-4 contains a faint band. This faint band might be indicative of a few colonies that show up on NA and MRS-agar plates, even after aseptic handling (data not shown). This signal might get picked up due to saturating PCR conditions. As we are not bleaching assay flies or using any antibiotics to avoid their toxic effects, we are essentially maintaining flies without any chemicals for about two consecutive generations, i.e., for ∼30 days (Supplementary Figure S4). So, there might be a tradeoff between the efficiency of keeping all the microbes away without any chemicals and how long that efficiency can be sustained without any chemical agent. Another possibility is that the signal is from very few non-culturable bacteria.

These results match the observations in Supplementary Text S5.1.1 obtained after plating flies on NA and MRS. Together, the evidence showed that both MB-MBL populations in a with-microbe environment harbor microbiome and in microbe-free environments have extremely few to no microbes.

**Figure S8.**
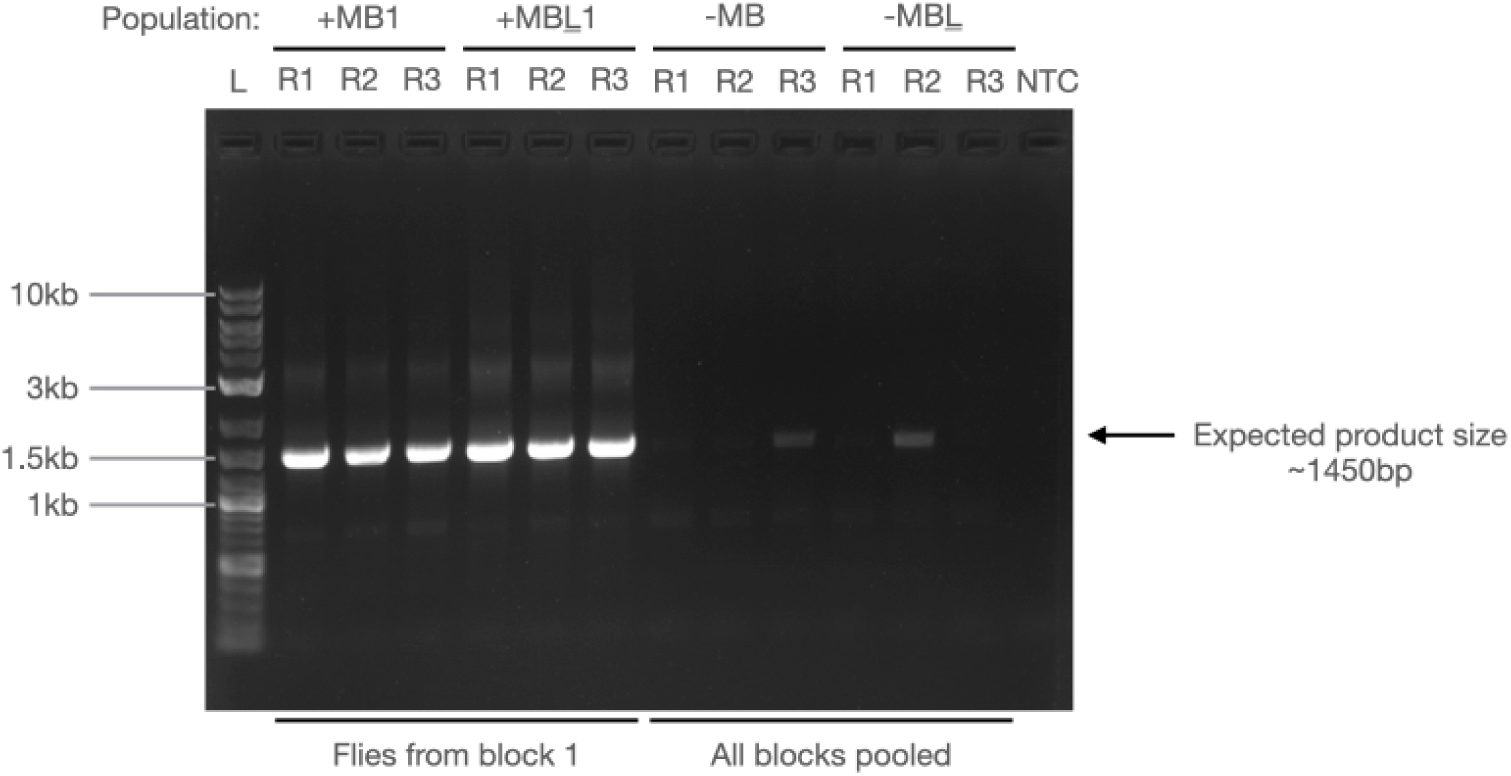
PCR on MB1 and MBL1-4 to confirm the status of microbiome.

#### S5.2 Validation of microbiome composition of flies raised in with-microbes assay environment

##### S5.2.1 MB-MBL generation 17-20 assays

In the previous section, through plating fly homogenates and PCR, we showed that flies raised in the with-microbes environment indeed have the microbes with them. However, the PCR only indicates that microbes are present - it does not tell us the composition of the reconstituted microbiome. In Supplementary Figure S7, we have established that the number of microbes received by MB and MBL populations is similar. In this section, we test if the reconstituted microbiome composition is similar for MB-MBLs, and this, in turn, matches the source microbiome composition of NDB1.

To validate that the composition of the reconstituted microbiome of all four blocks (MB1-4 and MBL1-4) reflects the native NDB1 composition, we performed amplicon sequencing on the V3-V4 region of the 16S rRNA gene (Supplementary Figure S9, method details in Supplementary Text S1.4). Three biological replicates were processed per population, giving us 24 samples (8 populations: 4 MBs + 4 MBLs, 3 replicates each). The native source pool of NDB1 (at Gen 56) was also sequenced.

Thus, Supplementary Figure S9 shows a total of 27 samples (24 from MB1-4-MBL1-4 and 3 from NDB1).

The MB1-4-MBL1-4 populations also mirror this broad microbiome composition of NDB1 with the dominance of *Acetobacter* spp. but with small fluctuations across MB-MBLs in *Acetobacter* and *Lactobacillus* fractions (Supplementary Figure S9). (In 2020, the old genus *Lactobacillus* was reclassified into 25 new different genera. *Lactiplantibacilus* and *Levilactobacilus* both belong to the old composite *Lactobacillus* genus.)

**Figure S9.**
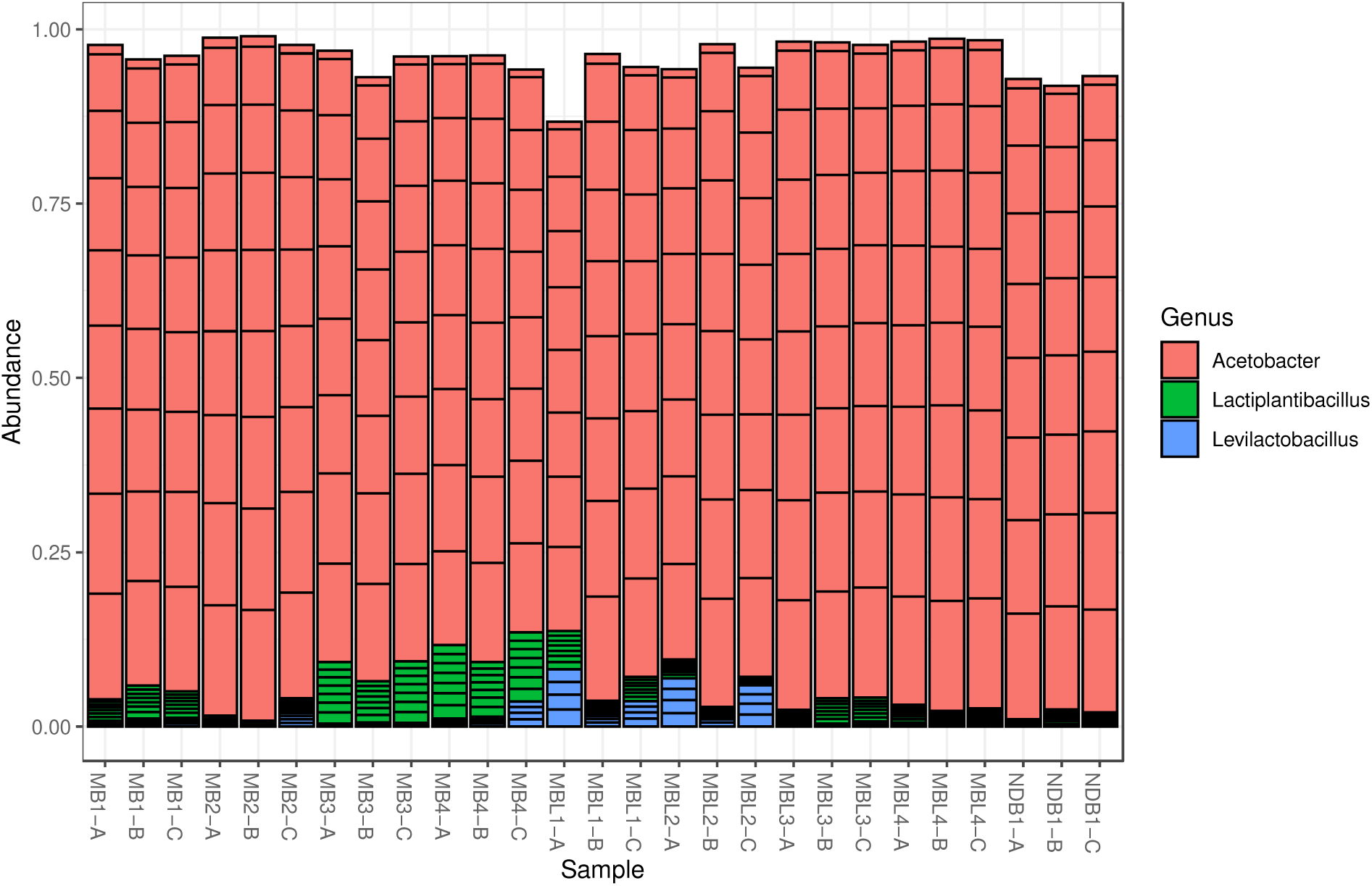
Validation of microbiome composition after reconstitution in Gen 17-20 assays. The result of population-level 16S rRNA amplicon sequencing across four replicate populations of MB1-4 and four replicate populations of MBL1-4. Native microbiome composition of source population NDB1 is also given for the reference (last three columns). Most abundant 20 taxa are shown for each population. For each population, three independent replicates where processed (labelled “A”, “B”, “C”). Each replicate consisted of pooled DNA extracted from surface sterilized full body homogenates of 8-10 females. The sequencing method and sequencing data analysis are described in Supplementary Text S1.4 and S1.5 respectively.

##### S5.2.2 MB-MBL generation 54-57 assays

In generation 54-57 assays, we sequenced microbiomes from MB1 and MBL1 that were raised in the with-microbes environments (Supplementary Figure S10). Together, the sequencing results for generations 17-20 and for generations 54-57 show that microbiome compositions of MBs and MBLs are broadly similar to each other and to the source microbiome of NDB1 (Supplementary Figure S9-last three lanes).

**Figure S10.**
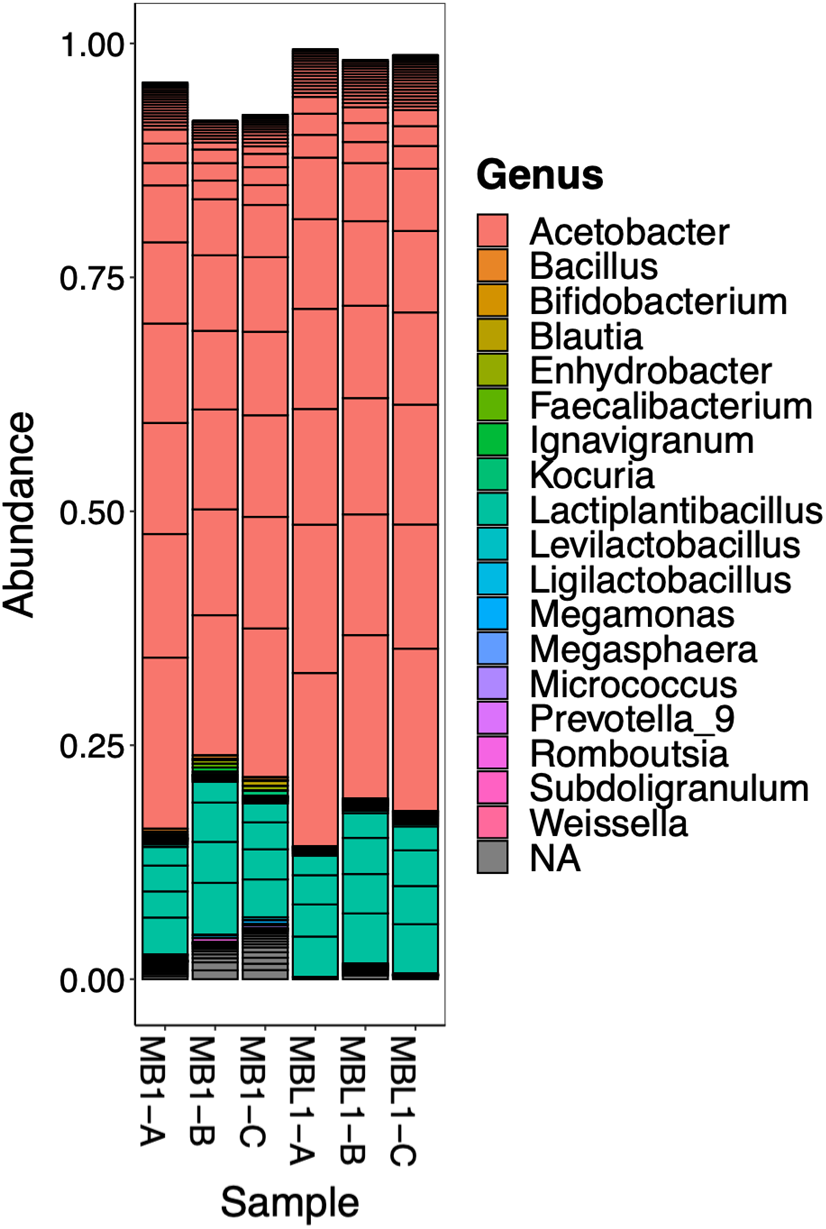
Validation of microbiome composition after reconstitution in Gen 54-57 assays. Most abundant 100 taxa are shown for each population. For each population, three independent replicates where processed (labelled “A”, “B”, “C”). Each replicate consisted of pooled DNA extracted from surface sterilized full body homogenates of 8-10 females. The sequencing method and sequencing data analysis are described in Supplementary Text S1.4 and S1.5 respectively.

### Text S6 - First phenotypic assessment of experimental evolution lines at host generations 17-20

#### S6.1 Male traits in host generations 17-20

None of the four phenotypes tested for males – adult body weight, egg-to-adult development time, desiccation resistance, and locomotor activity- showed an effect of selection as assessed by the paired t-test over block means (Supplementary Figure S11A-D). Supplementary Tables S5 and S6 provide details of the statistical comparison for the with-microbe assay environment and microbe-free assay environment, respectively.

**Figure S11.**
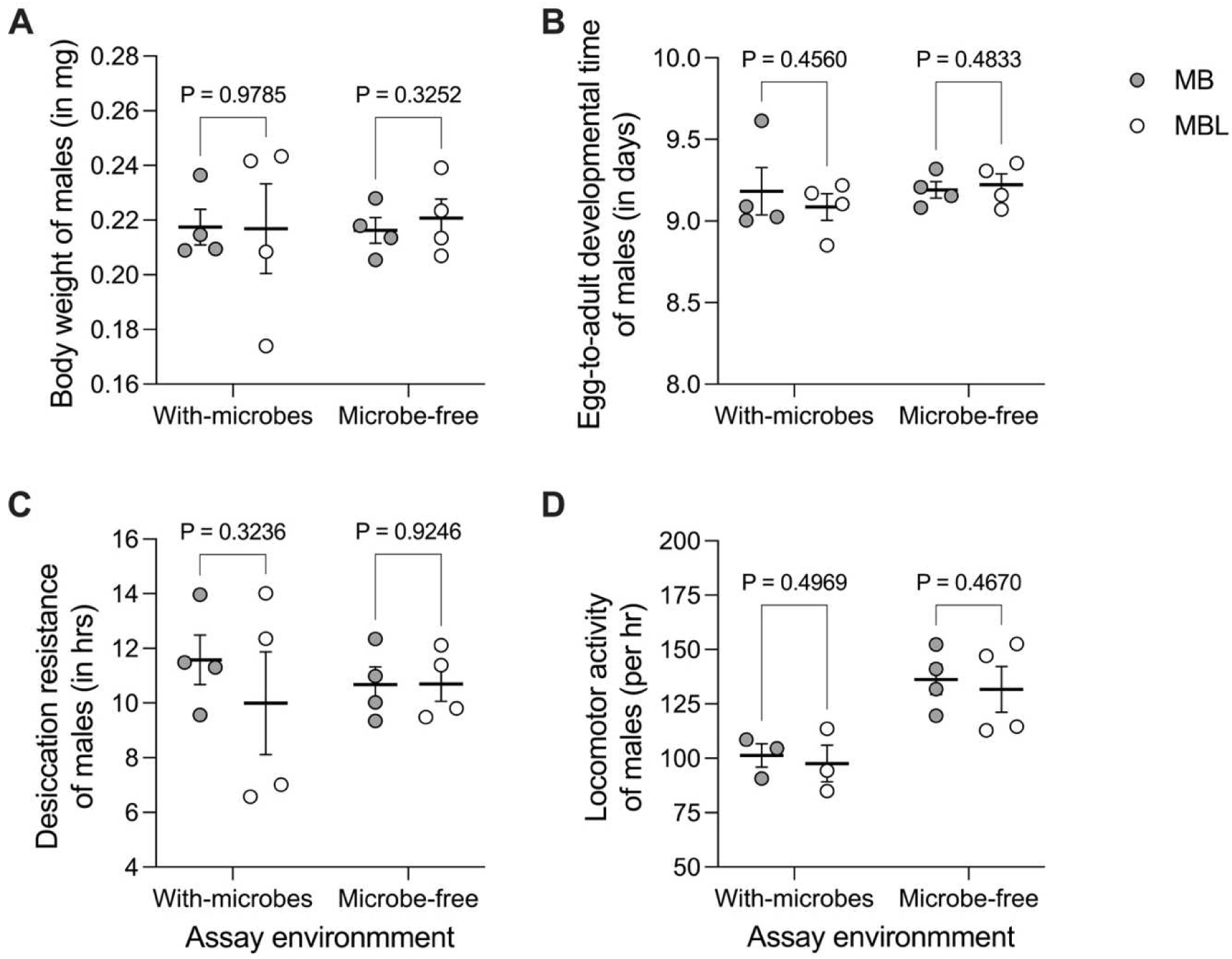
Male traits assessed in Gen 17-20 in with-microbe and microbe-free environments. (A) Body weight. (B) Egg-to-adult development time. (C) Desiccation resistance. (D) Locomotor activity using DAM system. In each assay environment, four filled circles are means for each of the four control populations MB1-4 and, similarly, four empty circles are means for each of the four selected populations MBL1-4. The black horizontal lines represent the grand means over four population means and the error bars represent the SEM. For each host phenotype, the grand mean over MB1-4 was compared against the grand mean over MBL1-4 using a paired t-test. DAM data for block 4 in with-microbes environment is not included here in the analysis.

#### S6.2 Female traits in host generations 17-20

In the case of females, egg-to-adult development time, desiccation resistance, and locomotor activity didn’t differ between MBs and MBLs (Supplementary Figure S12A-D) (Supplementary Tables S5 and S6). The only trait in females that showed some difference between the MBs and the MBLs was female body weight in the microbe-free environment (Paired t-test, t3 = 3.05, P = 0.0554, Cohen’s d = 0.81 (large), Supplementary Figure S12A). Here, although the p-value is slightly larger than our pre-determined level of statistical significance (i.e., 0.05), we still interpret this trait as the effect size was found to be large.

**Figure S12.**
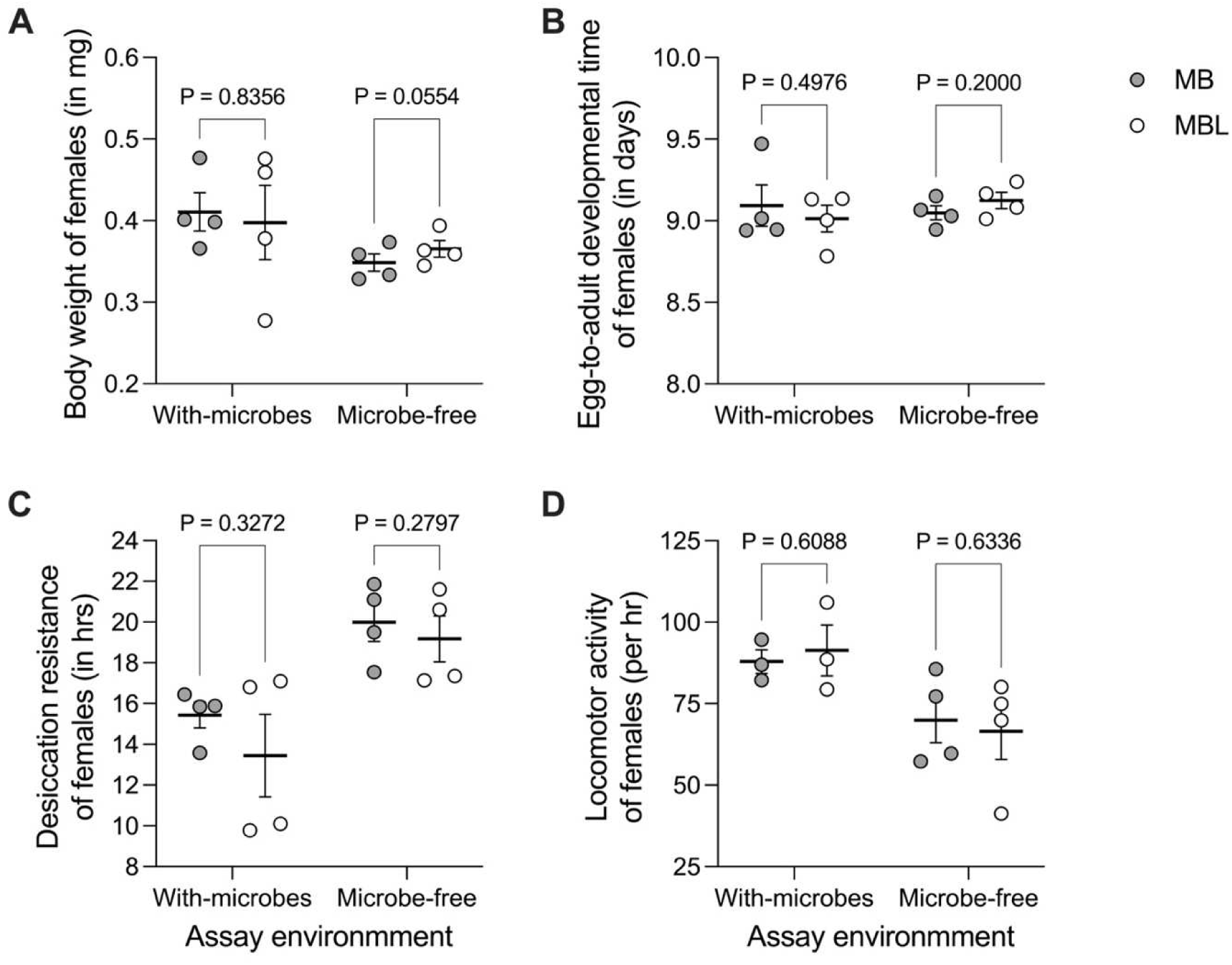
Female traits assessed in Gen 17-20 in with-microbe and microbe-free environments. (A) Body weight. (B) Egg-to-adult development time. (C) Desiccation resistance. (D) Locomotor activity using DAM system. In each assay environment, four filled circles are means for each of the four control populations MB1-4 and similarly, four empty circles are means for each of the four selected populations MBL1-4. The black horizontal lines represent the grand means over four population means and the error bars represent the SEM. For each host phenotype, the grand mean over MB1-4 was compared against the grand mean over MBL1-4 using a paired t-test. DAM data for block 4 in with-microbes environment is not included here in the analysis.

#### S6.3 Pair traits in host generations 17-20

In traits where both males and females were involved in the assays - percent survival to adulthood, female fecundity over 12 hrs (assayed in the presence of a male), and mating latency - did not show any difference (Supplementary Figure S13A-C, Supplementary Tables S5 and S6). The mating time for MBL1-4 was lower than MB1-4 in the with-microbes environment (Paired t-test, t3 = 3.67, P = 0.0349, Cohen’s d = 2.35 (large), Supplementary Figure S13D).

**Figure S13.**
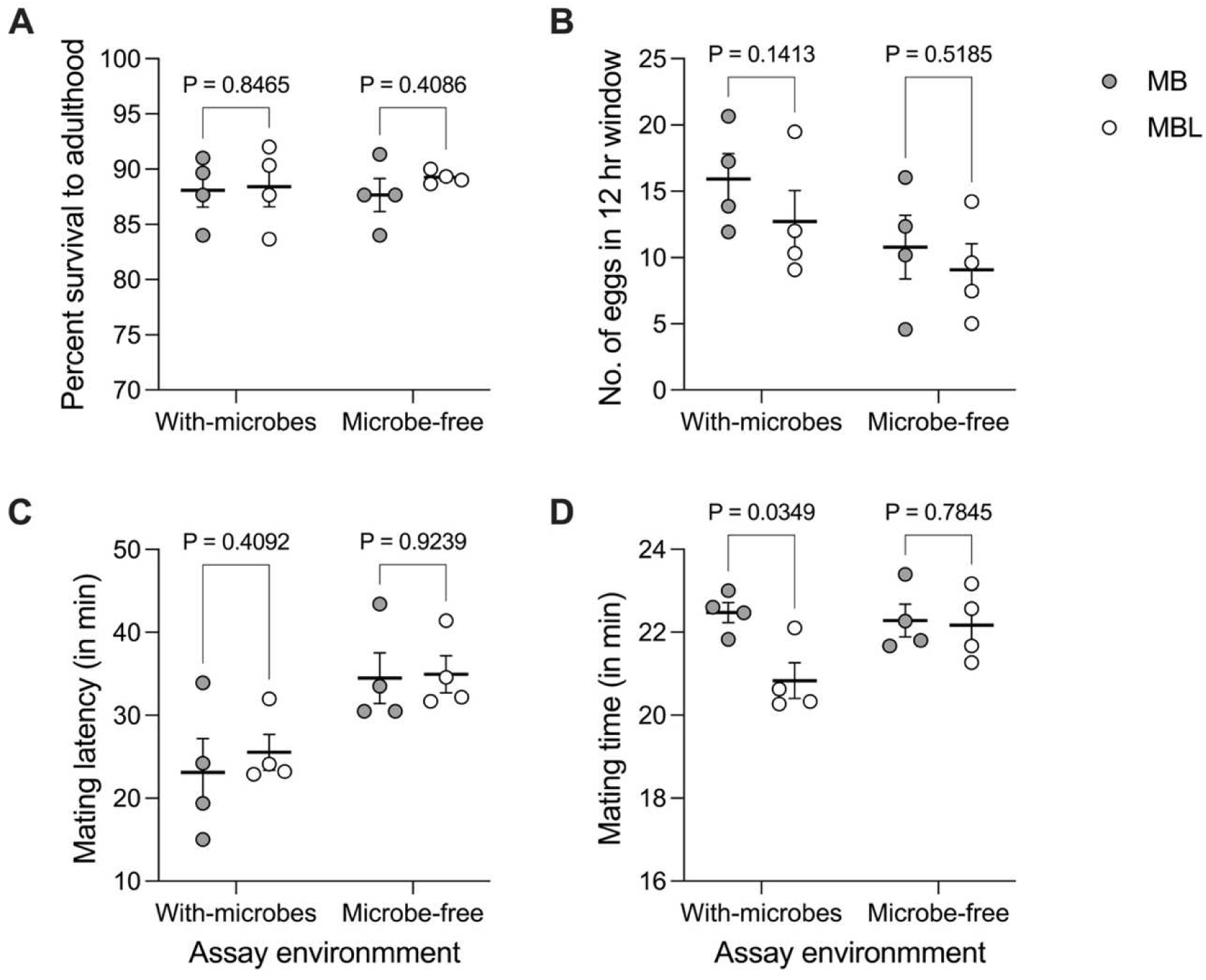
Pair traits assessed in Gen 17-20 in with-microbe and microbe-free environments. (A) Survival to adulthood. (B) Female fecundity. (C) Mating latency. (D) Mating time. In each assay environment, four filled circles are means for each of the four control populations MB1-4 and similarly, four empty circles are means for each of the four selected populations MBL1-4. The black horizontal lines represent the grand means over four population means and the error bars represent the SEM. For each host phenotype, the grand mean over MB1-4 was compared against the grand mean over MBL1-4 using a paired t-test.

**Table S5.** Summary of all assays in the <u>with-microbes environment (Gen 17-20).</u> T refers to the grand mean of the MBLs, while C refers to the grand mean of the MBs.

|  | Assay | Test statistic | P-value | % change (T - C)/C | Cohen's d | Effect size* | Sample size<sup>s</sup> |
| --- | --- | --- | --- | --- | --- | --- | --- |
| 1 | Dry body weight (Male) | $t_3 = 0.03$ | 0.9785 | -0.3 | 0.02 | Small | 10 |
| 2 | Dry body weight (Female) | $t_3 = 0.23$ | 0.8356 | -3.1 | 0.18 | Small | 10 |
| 3 | Development time (Males) | $t_3 = 0.85$ | 0.4560 | -1.1 | 0.41 | Small | 10 |
| 4 | Development time (Females) | $t_3 = 0.77$ | 0.4976 | -0.9 | 0.37 | Small | 10 |
| 5 | Survival to adulthood | $t_3 = 0.21$ | 0.8465 | 0.4 | 0.10 | Small | 10 |
| 6 | Fecundity (Females) | $t_3 = 1.96$ | 0.1413 | -20.1 | 0.75 | Medium | 50 |
| 7 | Desiccation resistance (Males) | $t_3 = 1.18$ | 0.3236 | -13.7 | 0.54 | Medium | 10 |
| 8 | Desiccation resistance (Females) | $t_3 = 1.17$ | 0.3272 | -12.9 | 0.66 | Medium | 10 |
| 9 | Locomotor activity (Males) | $t_3 = 0.82$ | 0.4969 | -3.7 | 0.30 | Small | 25 to 32 |
| 10 | Locomotor activity (Females) | $t_3 = 0.60$ | 0.6088 | 3.9 | 0.32 | Small | 31 to 32 |
| 11 | Mating Latency | $t_3 = 0.96$ | 0.4092 | 10.4 | 0.37 | Small | 30 |
| 12 | Mating time | $t_3 = 3.67$ | 0.0349 | -7.3 | 2.35 | Large | 30 |

**Table S6.**
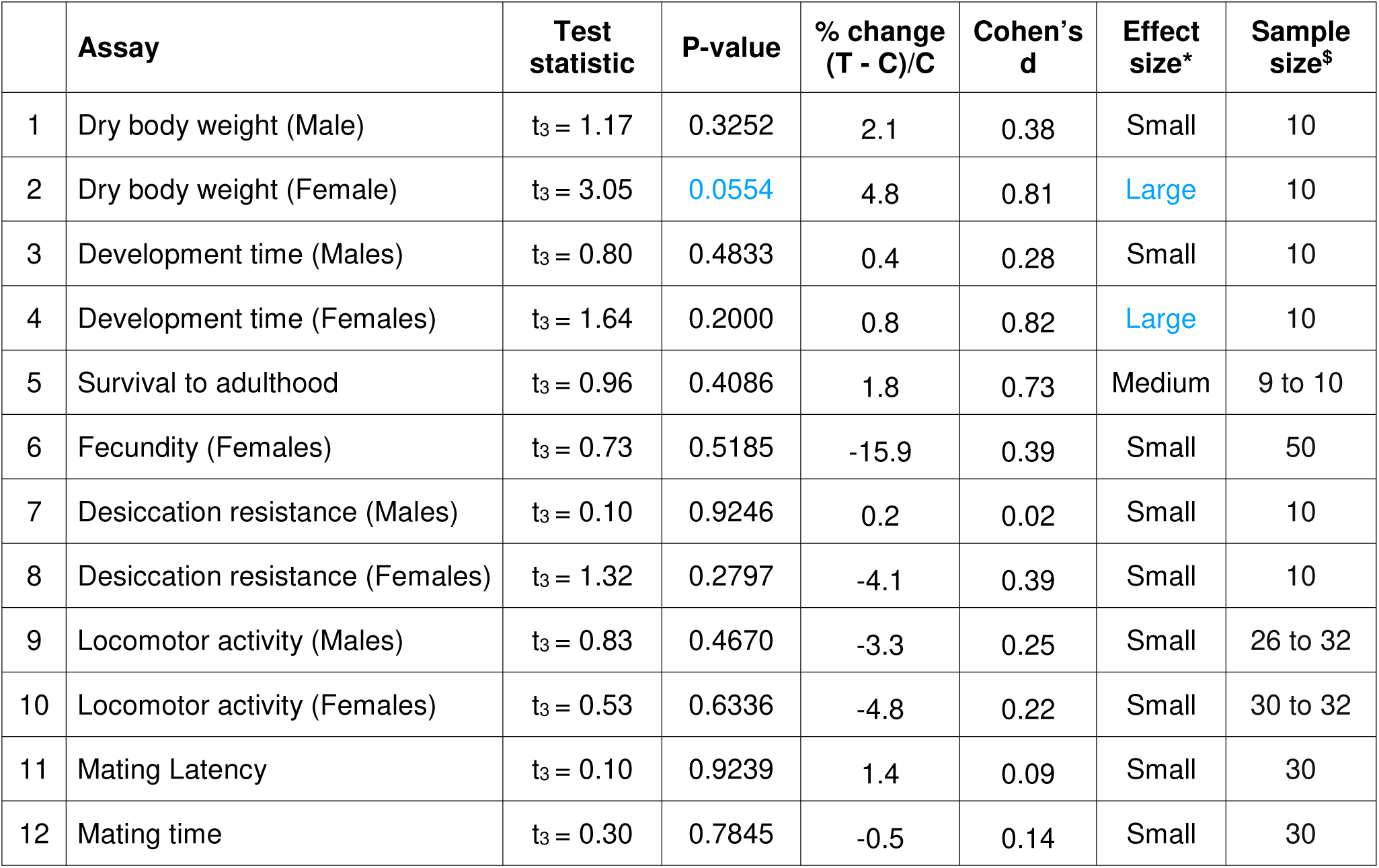
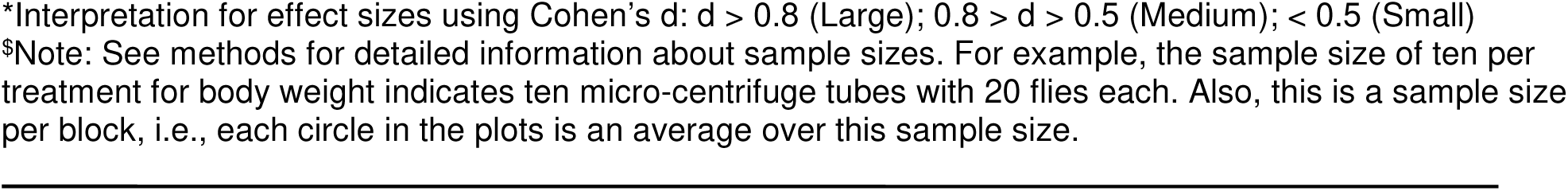
Summary of all assays in the <u>microbe-free environment (Gen 17-20).</u> T refers to the grand mean of the MBLs, while C refers to the grand mean of the MBs.

|  | Assay | Test statistic | P-value | % change (T - C)/C | Cohen's d | Effect size* | Sample size<sup>s</sup> |
| --- | --- | --- | --- | --- | --- | --- | --- |
| 1 | Dry body weight (Male) | $t_3 = 1.17$ | 0.3252 | 2.1 | 0.38 | Small | 10 |
| 2 | Dry body weight (Female) | $t_3 = 3.05$ | 0.0554 | 4.8 | 0.81 | Large | 10 |
| 3 | Development time (Males) | $t_3 = 0.80$ | 0.4833 | 0.4 | 0.28 | Small | 10 |
| 4 | Development time (Females) | $t_3 = 1.64$ | 0.2000 | 0.8 | 0.82 | Large | 10 |
| 5 | Survival to adulthood | $t_3 = 0.96$ | 0.4086 | 1.8 | 0.73 | Medium | 9 to 10 |
| 6 | Fecundity (Females) | $t_3 = 0.73$ | 0.5185 | -15.9 | 0.39 | Small | 50 |
| 7 | Desiccation resistance (Males) | $t_3 = 0.10$ | 0.9246 | 0.2 | 0.02 | Small | 10 |
| 8 | Desiccation resistance (Females) | $t_3 = 1.32$ | 0.2797 | -4.1 | 0.39 | Small | 10 |
| 9 | Locomotor activity (Males) | $t_3 = 0.83$ | 0.4670 | -3.3 | 0.25 | Small | 26 to 32 |
| 10 | Locomotor activity (Females) | $t_3 = 0.53$ | 0.6336 | -4.8 | 0.22 | Small | 30 to 32 |
| 11 | Mating Latency | $t_3 = 0.10$ | 0.9239 | 1.4 | 0.09 | Small | 30 |
| 12 | Mating time | $t_3 = 0.30$ | 0.7845 | -0.5 | 0.14 | Small | 30 |

\*Interpretation for effect sizes using Cohen's d: $d > 0.8$ (Large); $0.8 > d > 0.5$ (Medium); $< 0.5$ (Small) \$Note: See methods for detailed information about sample sizes. For example, the sample size of ten per treatment for body weight indicates ten micro-centrifuge tubes with 20 flies each. Also, this is a sample size per block, i.e., each circle in the plots is an average over this sample size.

### Text S7 – RNA-Seq result details

#### S7.1 No major changes in the gene expression profile of MB1 vs MBL1 in the microbe-free environment

In the microbe-free regime, where both selected and control populations were assayed without their microbiome, only a few transcripts showed differential expression when analyzed using *DESeq2* (Supplementary Figure S14). Only the significant genes with Log2(Fold Change) > 1.5 and adjusted P-value < 0.05 are highlighted in Figure S14. Except for nAChRbeta2, which is a nicotinic acetylcholine receptor subunit, and CG13617, which codes for cilium assembly protein DZIP1 (zinc finger transcription factor), the functions of the other genes (CG14625, CG12693) are not annotated in *Drosophila melanogaster*.

**Figure S14.**
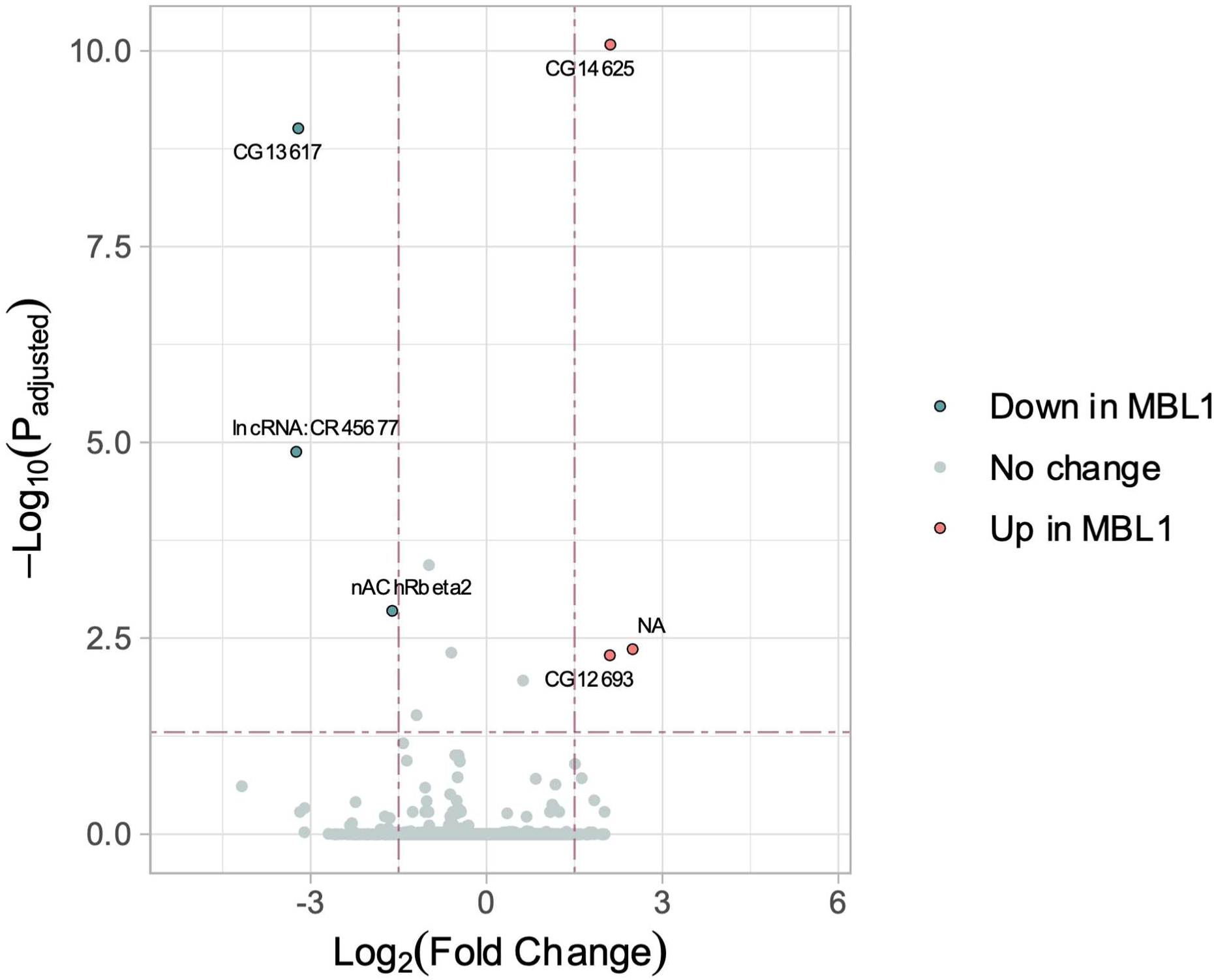
Differentially expressed genes in MBL1 vs MB1 in the no-microbe environment using DESeq2. The red dots represent the up-regulated genes, while blue dots represent down-regulated genes in MBL1. The protocol for the RNA-Seq is described in the method section of the main text.

#### S7.2 List of differentially regulated genes in with-microbe assay environment

**Table S7.** List of <u>up-regulated</u> genes in MBL1 in RNA-Seq (with-microbes environment)

| No. | Gene ID | Log<sub>2</sub>(FC) | Adjusted P-value | Symbol | Chr | Description |
| --- | --- | --- | --- | --- | --- | --- |
| 1 | FBgn0036948 | 2.59 | 5.29E-46 | CG7298 | 3L | - |
| 2 | FBgn0041579 | 2.74 | 8.56E-32 | AttC | 2R | Attacin-C |
| 3 | FBgn0010388 | 3.43 | 4.99E-31 | Dro | 2R | Drosocin |
| 4 | FBgn0034407 | 2.47 | 1.91E-10 | DptB | 2R | Diptericin B |
| 5 | FBgn0286723 | 2.63 | 2.44E-10 | hll | 2L | heimdall |
| 6 | FBgn0014865 | 2.28 | 1.46E-07 | Mtk | 2R | Metchnikowin |
| 7 | FBgn0004240 | 3.11 | 1.55E-05 | DptA | 2R | Diptericin A |
| 8 | FBgn0052284 | 2.28 | 0.0004 | CG32284 | 3L | - |
| 9 | FBgn0034957 | 3.78 | 0.0039 | Rsph4a | 2R | Radial spoke head protein 4a |
| 10 | FBgn0034712 | 1.76 | 0.0122 | Alp8 | 2R | Alkaline phosphatase 8 |
| 11 | FBgn0037323 | 2.37 | 0.0227 | CG2663 | 3R | - |
| 12 | FBgn0012042 | 1.64 | 0.0490 | AttA | 2R | Attacin-A |

**Table S8.** List of <u>down-regulated</u> genes in MBL1 in RNA-Seq (with-microbes environment)

| No. | Gene ID | Log<sub>2</sub>(FC) | Adjusted P-value | Symbol | Chr | Description |
| --- | --- | --- | --- | --- | --- | --- |
| 1 | FBgn0033978 | -1.53 | 3.30E-19 | Cyp6a23 | 2R | Cyp6a23 |
| 2 | FBgn0001225 | -3.94 | 1.75E-13 | Hsp26 | 3L | Heat shock protein 26 |
| 3 | FBgn0001229 | -2.03 | 1.43E-09 | Hsp67Bc | 3L | Heat shock gene 67Bc |
| 4 | FBgn0013279 | -2.37 | 7.09E-07 | Hsp70Bc | 3R | Heat-shock-protein-70Bc |
| 5 | FBgn0001224 | -2.74 | 5.69E-06 | Hsp23 | 3L | Heat shock protein 23 |
| 6 | FBgn0039151 | -1.84 | 6.07E-06 | CG13607 | 3R | - |
| 7 | FBgn0013276 | -2.48 | 1.55E-05 | Hsp70Ab | 3R | Heat-shock-protein-70Ab |
| 8 | FBgn0051354 | -2.28 | 1.55E-05 | Hsp70Bbb | 3R | Hsp70Bbb |
| 9 | FBti0059721 | -4.34 | 2.33E-05 | NA | 3R | - |
| 10 | FBgn0001230 | -3.18 | 2.48E-05 | Hsp68 | 3R | Heat shock protein 68 |
| 11 | FBti0019359 | -2.89 | 4.19E-05 | NA | 3R | - |
| 12 | FBgn0013278 | -2.50 | 0.0003 | Hsp70Bb | 3R | Heat-shock-protein-70Bb |
| 13 | FBgn0030769 | -1.75 | 0.0005 | CG13012 | X | - |
| 14 | FBgn0039152 | -1.60 | 0.0005 | Root | 3R | Rootletin |
| 15 | FBgn0036363 | -6.29 | 0.0006 | CG10140 | 3L | - |
| 16 | FBgn0013275 | -2.24 | 0.0009 | Hsp70Aa | 3R | Heat-shock-protein-70Aa |
| 17 | FBgn0013277 | -2.41 | 0.0023 | Hsp70Ba | 3R | Heat-shock-protein-70Ba |
| 18 | FBgn0052495 | -1.61 | 0.0078 | Gss2 | X | Glutathione synthetase 2 |
| 19 | FBgn0039307 | -2.28 | 0.0079 | CR13656 | 3R | - |
| 20 | FBti0020330 | -2.39 | 0.0227 | NA | 3R | - |
| 21 | RR43222_transposable_element | -2.87 | 0.0300 | NA | X | - |
| 22 | FBti0059720 | -2.94 | 0.0306 | NA | 3R | - |
| 23 | FBti0019362 | -4.05 | 0.0393 | NA | 3R | - |
| 24 | FBgn0034784 | -2.69 | 0.0476 | CG9826 | 2R | - |

#### S7.3 GO analysis of differentially regulated genes in the with-microbe assay environment

To see the biological processes in which these genes are involved, we used the ShinyGo tool (version 0.80) (http://bioinformatics.sdstate.edu/go/) (Ge et al., 2020) for gene ontology enrichment analysis with ‘GO biological process’ pathway database. We found that the upregulated genes belong to the pathway involved in the fly’s humoral immune response (Figure S15), while down-regulated genes belong to the stress response pathway (Figure S16).

**Figure S15.**
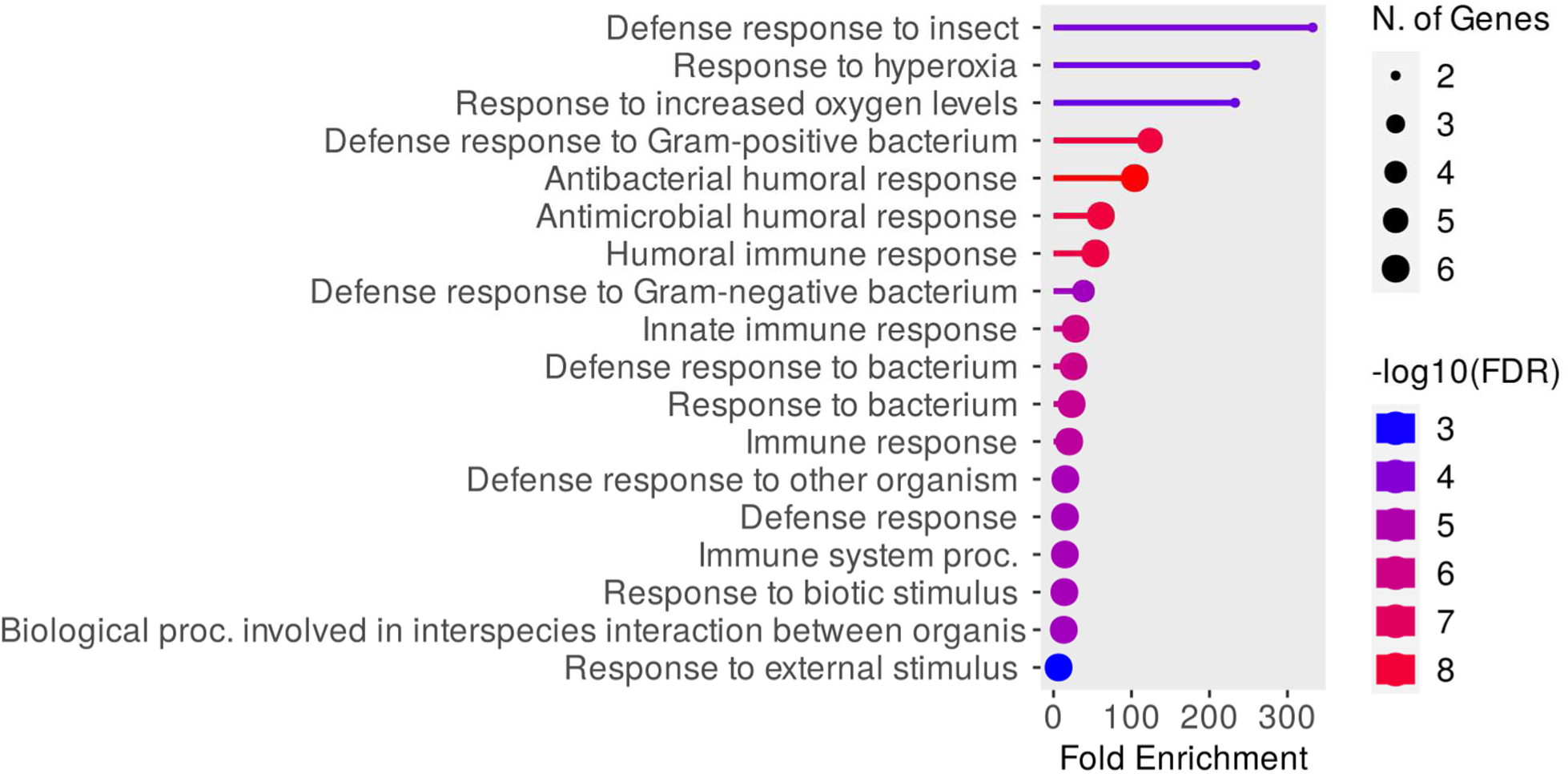
Gene ontology (GO) enrichment analysis for 12 Up-regulated genes.

**Figure S16.**
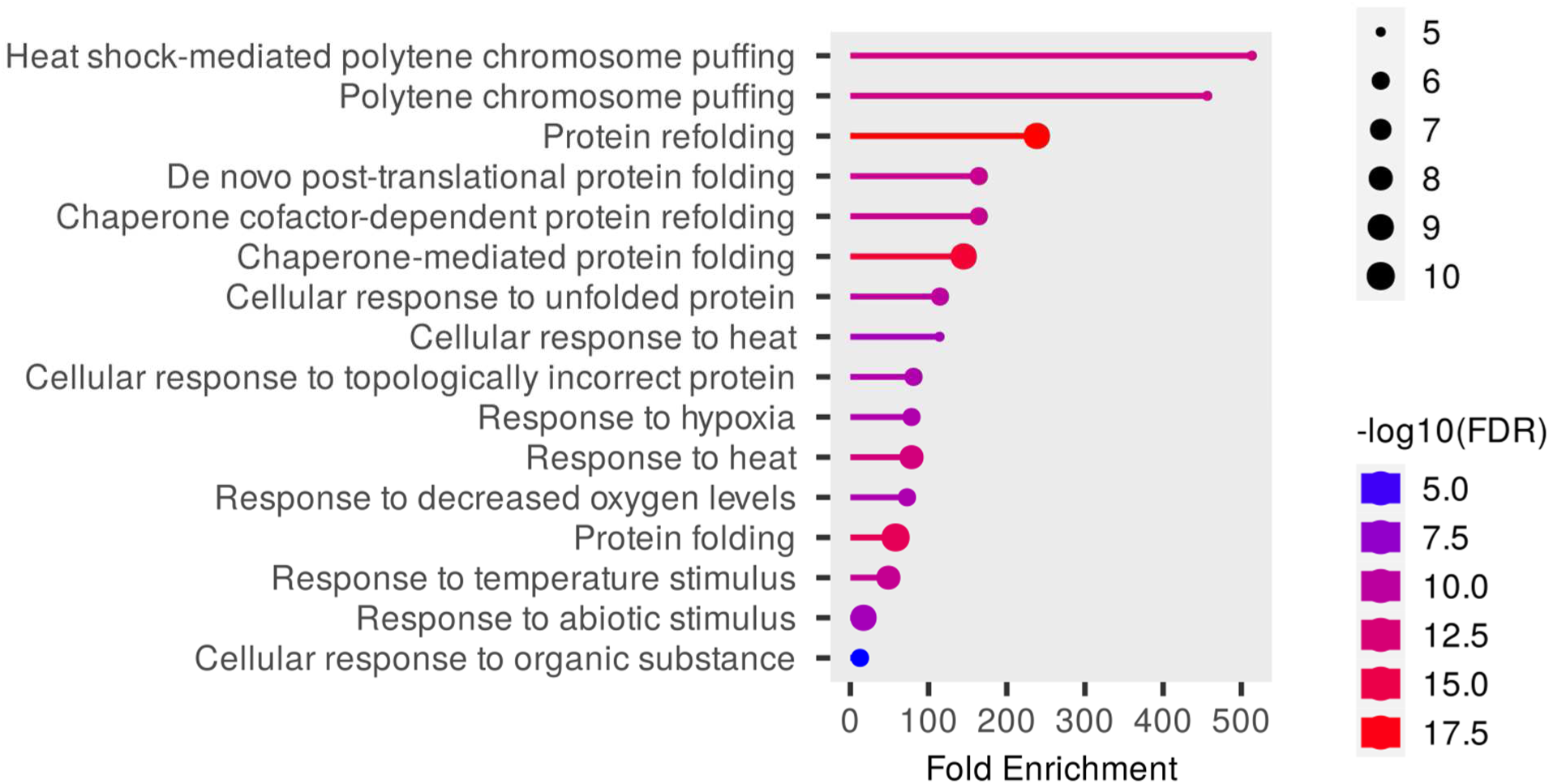
Gene ontology (GO) enrichment analysis for 24 down-regulated genes.

### Text S8 - Another set of differently expressed genes hints at the peritrophic matrix (PM) as a potential region of interest in the MBLs

Along with HSPs and AMPs, another set of genes that have an altered expression are chitin-interacting genes. CG7298 and CG32284 are up-regulated chitin-binding enzymes in MBL1, and down-regulated CG10140 is a probable chitinase. As the RNA-seq is done on the full body of flies, it is difficult to predict what this means for the fly. But, the annotation of CG7298 in FlyBase offers a clue. CG7298 is expressed in the proventriculus, which is part of the fly digestive system. A recent study supports the notion that proventriculus is the site where gut bacteria stably interact with flies, possibly through interactions with N-acetyl glucosamine, which is a building block of chitin (Dodge et al., 2023). Moreover, *Drosophila melanogaster* CG7298 has an ortholog called peritrophin-48 in related *Drosophila* species, which is a component of the peritrophic matrix. The peritrophic matrix is a chitinous layer that separates food bolus from the gut epithelia. This is also implicated as a physical barrier that keeps gut bacteria away from the gut tissue (Charroux & Royet, 2012; Erlandson et al., 2019; Kuraishi et al., 2011; Rodgers et al., 2017). Taken together, we think this is the region where chitin-interacting proteins are carrying out various processes that might be of interest. However, the limited available information about these hits prevents us from drawing any further conclusions at this point in time. In the future, we plan to confirm all the RNA-seq hits and do additional experiments that can give us more insights.

In a recent study, Kathedar et al. showed that the nicotinic acetylcholine receptor (nAchR) regulates gut barrier function in *Drosophila* through the remodeling of the peritrophic matrix (Katheder et al., 2023). We have seen that in our no-microbe assay regime, nAchRbeta2 gene expression showed down-regulation in MBL1. This raises the possibility that the processes of interest happening at the proventriculus that involve the peritrophic matrix might be mediated through nAch signaling.

